# Target Identification and Mechanistic Characterization of Indole Terpenoid Mimics: Proper Spindle Microtubule Assembly Is Essential for Cdh1-Mediated Proteolysis of CENP-A

**DOI:** 10.1101/2023.04.25.538200

**Authors:** Yan Peng, Yumeng Zhang, Ruan Fang, Hao Jiang, Gongcai Lan, Zhou Xu, Yajie Liu, Zhaoyang Nie, Lu Ren, Fengcan Wang, Shou-De Zhang, Yuyong Ma, Peng Yang, Hong-Hua Ge, Wei-Dong Zhang, Cheng Luo, Ang Li, Weiwei He

## Abstract

Centromere protein A (CENP-A), a histone H3 variant specific to centromeres, is crucial for kinetochore positioning and chromosome segregation. However, its regulatory mechanism in human cells remains incompletely understood. We conducted a structure-activity relationship (SAR) study of the cell cycle-arresting indole terpenoid mimic JP18 and found two more potent analogues, (+)-6-Br-JP18 and (+)-6-Cl-JP18. Tubulin was identified as a potential cellular target of these halogenated analogues by using the drug affinity responsive target stability (DARTS) based method. X-ray crystallography analysis revealed that both molecules bind to the colchicine-binding site of β-tubulin. Furthermore, we discovered that treatment of human cells with microtubule-targeting agents (MTAs), including these two compounds, led to CENP-A accumulation by destabilizing Cdh1, a co-activator of the APC/C E3 ubiquitin ligase. Our study establishes a link between microtubule dynamics and CENP-A accumulation using small-molecule tools and highlights the role of Cdh1 in CENP-A proteolysis.

## ▪ INTRODUCTION

Mitosis is a process in which a eukaryotic cell divides into two genetically identical cells, each containing approximately equal proportions of cellular components.[1,2] The mitotic spindle, a dynamic microtubule-based structure that forms during mitosis, is responsible for chromosome segregation.[3] Therefore, proper microtubule assembly and disassembly are crucial for mitosis.[4] Disrupting microtubule dynamics is an effective strategy for suppressing tumor cell proliferation.[5] Numerous microtubule-targeting agents (MTAs), such as paclitaxel and vincristine among natural products, and ixabepilone and eribulin among natural product analogues, are employed in clinical practice.[6] However, prolonged use of MTAs can lead to chemoresistance,[7,8] presumably due to defects in mitotic fidelity such as aneuploidy and chromosomal instability (CIN),[9–13] although the exact mechanism remains unclear.

Centromere protein A (CENP-A) is a histone H3 variant playing a key role in faithful chromosome segregation.[14,15] The centromere is a constricted region of the chromosome that links sister chromatids.[16] CENP-A replaces histone H3 in the histone octamer at the centromere, facilitating the formation of the centromere-specific nucleosome.[14,15] This nucleosome acts as a platform for the assembly of the kinetochore, which attaches the chromosome to the spindle microtubules.[17,18] Therefore, accurate chromosome segregation relies on proper CENP-A deposition at the centromere. Overexpression of CENP-A results in its mislocalization to non-centromere regions and the formation of ectopic kinetochores on chromosome arms, ultimately leading to segregation defects, aneuploidy, and CIN.[15,19,20] Notably, various human tumors exhibit abnormally high CENP-A levels, which significantly correlate with malignant progression and poor patient survival.[21–24]

CENP-A stability and centromeric deposition are modulated by ubiquitination across species.[25,26] Recently, there has been increasing attention on its regulation in human cells. Wang et al. discovered that the CUL4−DDB1−DCAF11 E3 ubiquitin ligase mediates the Ser68 phosphorylation-dependent poly-ubiquitylation of CENP-A, leading to its degradation in human cervical cancer cells (HeLa).[27] In addition, Niikura et al. reported that the mono-ubiquitylation of CENP-A at Lys124, promoted by the CUL4A-RBX1-COPS8 E3 ubiquitin ligase, is essential for CENP-A deposition at the centromere in HeLa cells.[28] However, the link between impaired microtubule dynamics and defective CENP-A regulation remains largely unexplored. Understanding this connection may shed light on the mechanistic basis of MTA-induced tumor evolution, including the emergence of acquired chemoresistance.

Indole terpenoids are a large class of natural products characterized by a hybrid structure consisting of a terpene moiety and an indole moiety.[29,30] They exhibit a wide range of biological activities, with increasing interest in their anticancer properties.[31,32] For instance, the penitrem-type indole diterpenoids have been shown to suppress the proliferation, migration, and invasion of human breast cancer cells.[33–35] Leveraging our expertise in indole terpenoid synthesis,[36–43] we have embarked on a program to identify anticancer agents among synthetic indole terpenoids and their analogues and to elucidate their mechanisms of action. In our previous study, the indole terpenoid mimic JP18 was found to induce cell cycle arrest in the G2/M phase and inhibit cancer cell proliferation.[44] Herein, we report the discovery of two halogenated analogues of JP18, (+)-6-Br-JP18 and (+)-6-Cl-JP18, as more potent cell cycle blockers through a structure−activity relationship (SAR) study. Label-free target identification based on drug affinity responsive target stability (DARTS) indicated tubulin as a potential cellular target of these compounds, and X-ray crystallography analysis confirmed their interaction with the colchicine-binding site of β-tubulin. Disrupting spindle microtubule dynamics by MTAs, including (+)-6-Br-JP18 and (+)-6-Cl-JP18, downregulates Cdh1, a co-activator of APC/C E3 ubiquitin ligase, leading to accumulation of its substrate CENP-A in human cells.

## ▪ RESULTS

### Identification of two potent cell cycle inhibitors by an SAR study of JP18

Our previous demonstrated that the indole terpenoid mimic JP18 (1) inhibited cell proliferation and induced cell cycle arrest in the G2/M phase in human lung cancer cells (A549).[44] To gain insight into the SAR of JP18, we constructed a focused library consisting of a series of analogues of 1 (Figure 1A) via a conjugate addition strategy.[37,42,44] These racemic compounds were evaluated for their effects on cell cycle progression in HeLa cells by using flow cytometry. As observed in A549 cells, compound 1 arrested the cell cycle in the G2/M phase in HeLa cells in a dose-dependent manner (Figures 1B and S1). The importance of the carbonyl group was assessed by testing compounds 2 and 3 (Figure 1); the reduced potency highlighted the critical role of this functionality. The effects of indole substituents were then examined. The *N*1-alkylated analogue (4) was essentially inactive in the concentration range of 1.0 to 10 *μ*M, and the introduction of a bromine atom at C4, C5, and C7 significantly decreased potency (see the data for compounds 5–7). To our delight, 6-Br-JP18 (8) demonstrated considerably stronger activity than 1. We subsequently tested analogues with various C6 substituents. 6-Cl-JP18 (9) exhibited potency essentially equivalent to that of 8, and 10–12 also displayed enhanced activity compared to 1. In contrast, the C6-florinated analogue (13) showed reduced potency, and compounds 14–16 containing larger C6 substituents were largely inactive in the tested concentration range. Notably, compounds 8 and 9 maintained efficacy in the concentration range of 0.10 to 2.0 *μ*M (Figure S2), with measured half-maximal inhibitory concentrations (IC_50_) of 1.35 and 2.07 *μ*M, respectively, compared to 9.40 *μ*M for 1 (Figure S3).

**Figure 1.**
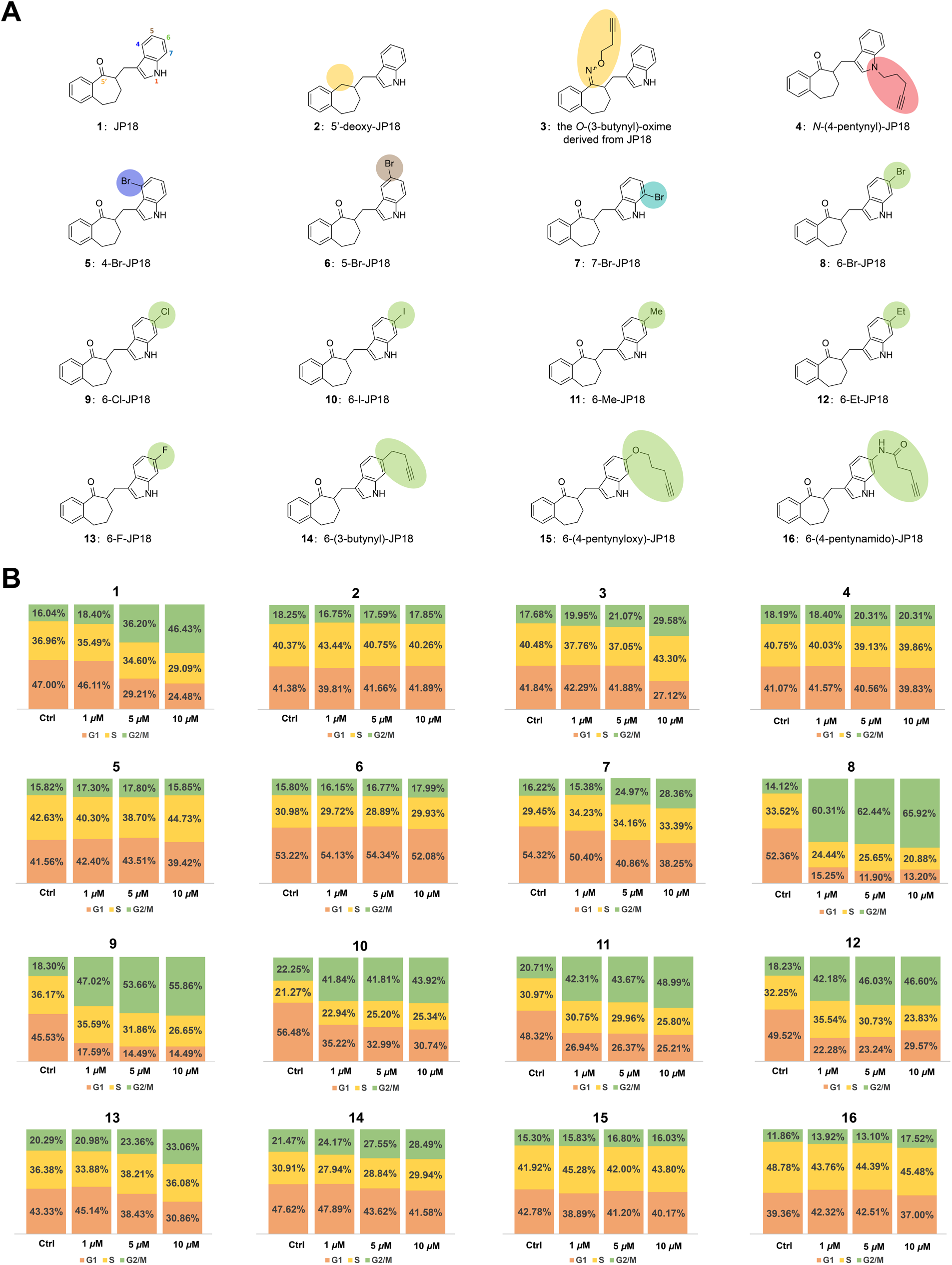
Cell cycle-arresting activity of JP18 and its analogues. (A) Structures of JP18 and its fifteen representative analogues. (B) Flow cytometry-based cell cycle analysis. HeLa cells were treated with the indicated compounds at specified concentrations for 8 h and then stained with propidium iodide (PI). DMSO was used as a control for the compounds. Bar graphs depict the percentages of cells in the G2/M (green), S (yellow), and G1 (orange) phases.

The enantiomeric forms responsible for the cell cycle-arresting activity of **8** and **9** were determined. Racemic samples of **8** and **9** were separated by high-performance liquid chromatography (HPLC) using a chiral stationary phase. Flow cytometry analysis revealed that both (+)-**8** and (+)-**9** induced cell cycle arrest in the G2/M phase in the concentration range of 0.10 to 2.0 *μ*M, whereas their enantiomers exhibited much weaker activity (Figures 2A–2D and S4). Furthermore, treatment of HeLa cells with both (+)-**8** and (+)-**9** increased cyclin B1 levels and decreased phospho-CDK1 (Tyr15) levels in a dose-and time-dependent manner (Figures 2E–2H), confirming their G2/M phase-arresting activity.[45] Notably, the absolution configuration of (+)-**8** has been elucidated by a combination of asymmetric synthesis[46] and computation, which will be reported elsewhere in due course.

**Figure 2.**
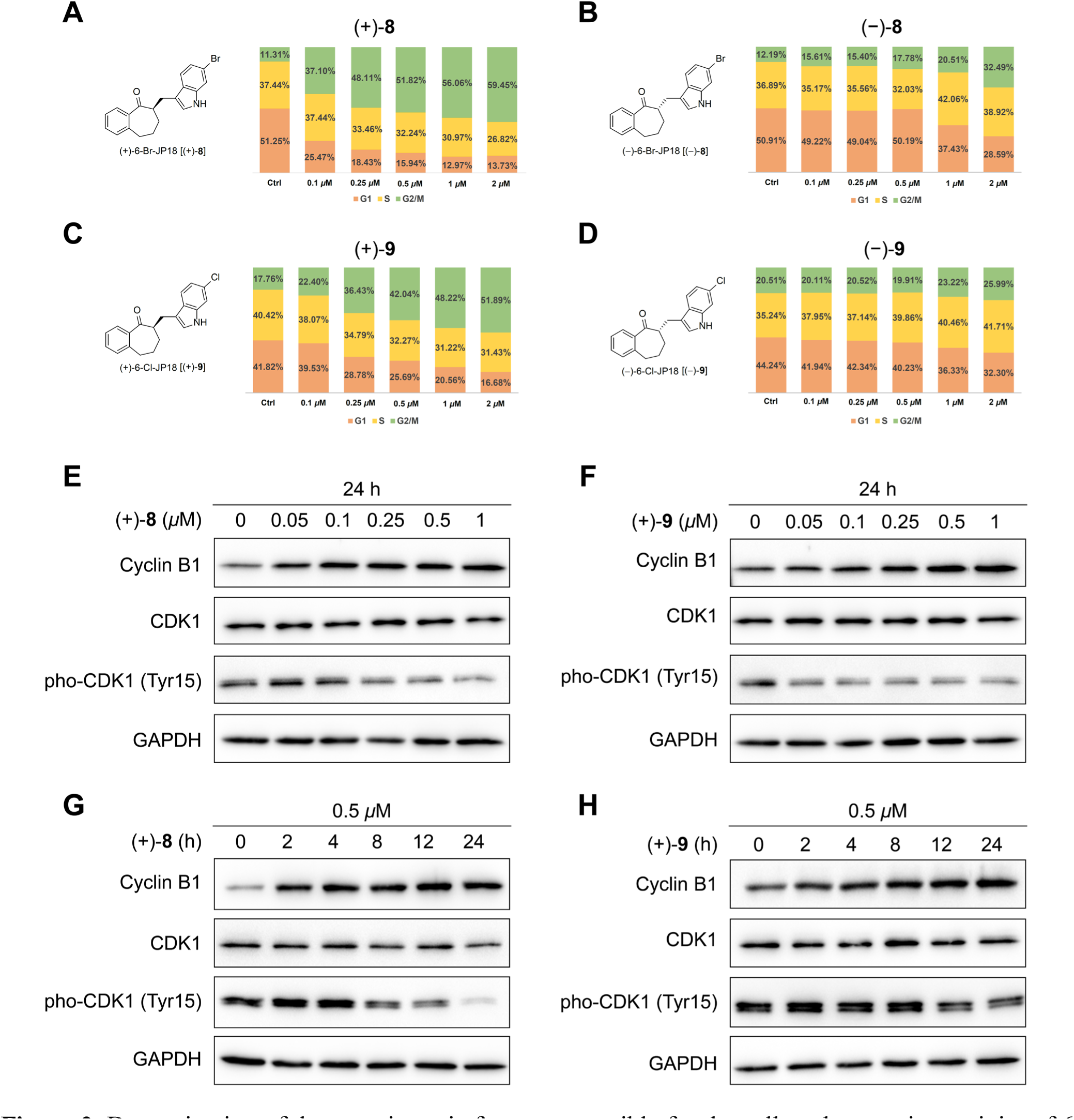
Determination of the enantiomeric forms responsible for the cell cycle-arresting activity of 6-Br-JP18 and 6-Cl-JP18. (A, B) The cell cycle-arresting effect of the two enantiomers of 6-Br-JP18 (**8**). HeLa cells were treated with (+)-**8** and (−)-**8**, respectively, at indicated concentrations for 8 h. (C, D) The cell cycle-arresting effect of the two enantiomers of 6-Cl-JP18 (**9**). HeLa cells were treated with (+)-**9** and (−)-**9**, respectively, at indicated concentrations for 8 h. Bar graphs depict the percentages of cells in the G2/M (green), S (yellow), and G1 (orange) phases. (E, F) Immunoblot analysis of G2/M phase markers in HeLa cells treated with (+)-**8** and (+)-**9**, respectively, at indicated concentrations for 24 h. (G, H) Immunoblot analysis of G2/M phase markers in HeLa cells treated with (+)-**8** and (+)-**9**, respectively, at 0.50 *μ*M for indicated durations. DMSO was used as a control for the compounds, and GAPDH was used as a loading control in immunoblotting.

### Compounds (+)-8 and (+)-9 disrupt spindle microtubule assembly and arrest cell cycle in the M phase

To elucidate their mode of action, we investigated whether (+)-**8** and (+)-**9** induce cell cycle arrest in the G2 phase or the M phase. As the flow cytometry method described above could not differentiate between the two phases, we examined key markers of mitosis. Phosphorylation of histone H3 at Ser10 and Ser28 is associated with chromosome condensation, which occurs from prophase to metaphase of mitosis.[47] Both (+)-**8** and (+)-**9** increased the levels of phosphorylated histone H3 in a dose- and time-dependent manner (Figures 3A–3D), indicating that they arrest the cell cycle in the M phase. Immunofluorescence analysis revealed that both compounds disrupted spindle microtubule assembly (Figure 3E). Furthermore, treatment with these compounds resulted in an uneven distribution of chromatids in the telophase of mitosis (Figures 3E8 and 3E12), which suggested a disruptive effect on centromere organization and function.

**Figure 3.**
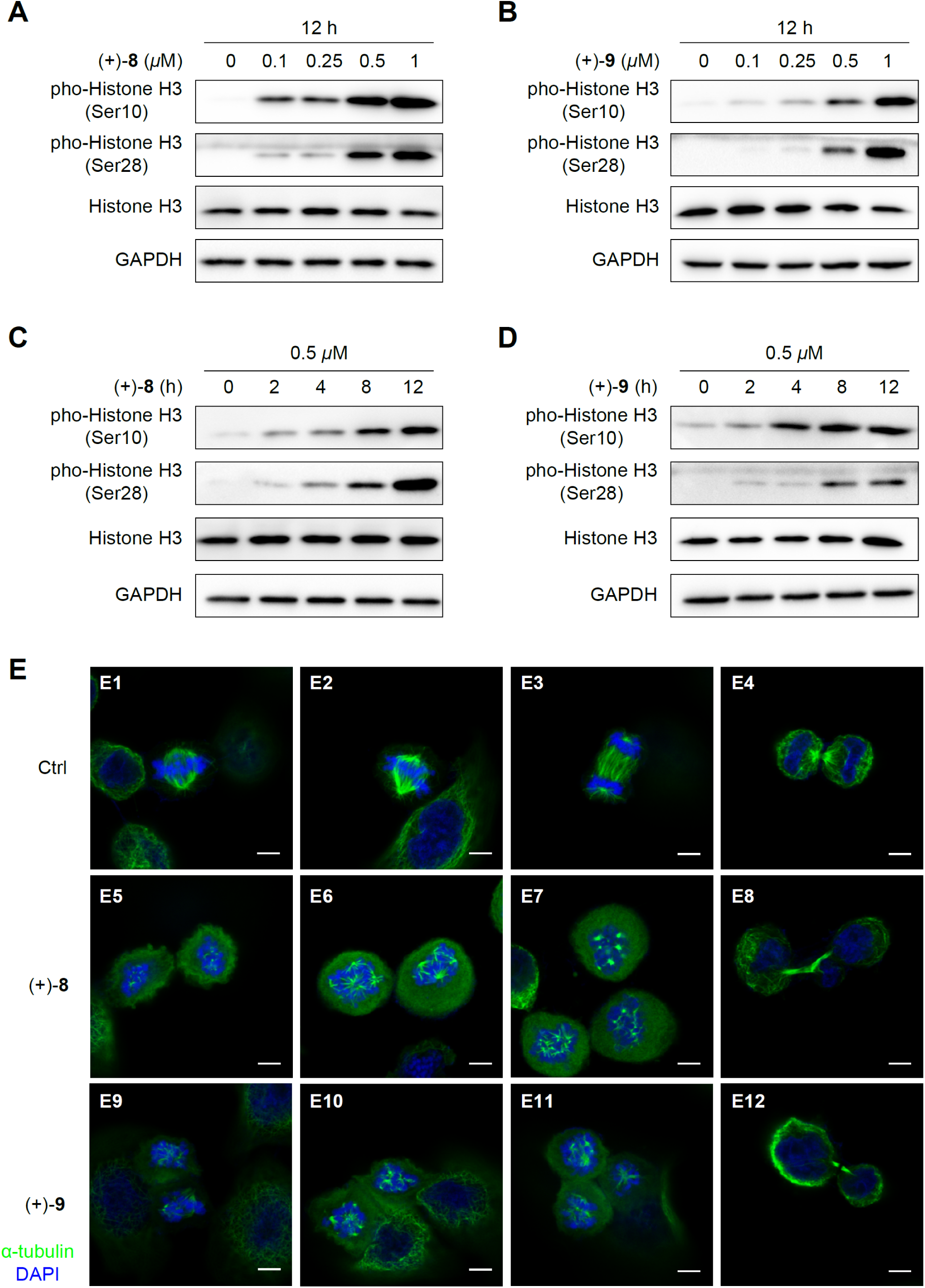
Effect of compounds (+)-**8** and (+)-**9** on M phase markers. (A, B) Immunoblot analysis of M phase markers in HeLa cells treated with (+)-**8** and (+)-**9**, respectively, at indicated concentrations for 12h. (C, D) Immunoblot analysis of M phase markers in HeLa cells treated with (+)-**8** and (+)-**9**, respectively, at 0.50 *μ*M for indicated durations. DMSO was used as a control for the compounds, and GAPDH was used as a loading control in immunoblotting. (E) Immunofluorescence analysis of spindle microtubules and chromosomes in HeLa cells treated with (+)-**8** and (+)-**9**, respectively, at 100 nM for 8 h. Cells were stained with the α-tubulin antibody (green) and DAPI (blue). Scale bars indicate 5.0 *μ*m.

Given their potent cytotoxicity, we examined the effect of (+)-**8** and (+)-**9** on a series of proteins associated with cell survival and death. The pro-survival protein B-cell lymphoma 2 (Bcl-2)[48] was downregulated in a dose- and time-dependent manner, while the DNA damage marker phospho-histone H2AX (Ser139) (γ-H2AX)[49] was upregulated (Figure S5). Additionally, increasing doses and durations of treatment with these compounds led to the escalation of poly(ADP-ribose) polymerase 1 (PARP1) cleavage (Figure S5), a widely used marker for apoptosis.[50] These results indicated that (+)-**8** and (+)-**9** induce DNA damage and apoptosis.

### Identification of α- and β-tubulins as potential cellular targets of compounds (+)-8 and (+)-9

The SAR study of JP18 highlighted the challenge in constructing a suitable small molecule probe for affinity-based target identification. Therefore, we resorted to the label-free method based on DARTS, which utilizes changes in the stability of the target protein against proteolysis due to interaction with the small molecule.[51,52] Among the proteins identified by mass spectrometry, α- and β-tubulins stood out as potential targets of (+)-**8** (Figures 4A and 4B), a conclusion supported by its disruption of spindle microtubule assembly (Figure 3E). To confirm the direct binding of (+)-**8** to tubulin, we conducted an in vitro tubulin polymerization assay. At a concentration of 5.0 *μ*M, compound (+)-**8** significantly inhibited tubulin polymerization, similar to the reference MTA nocodazole (Figure 4C).

**Figure 4.**
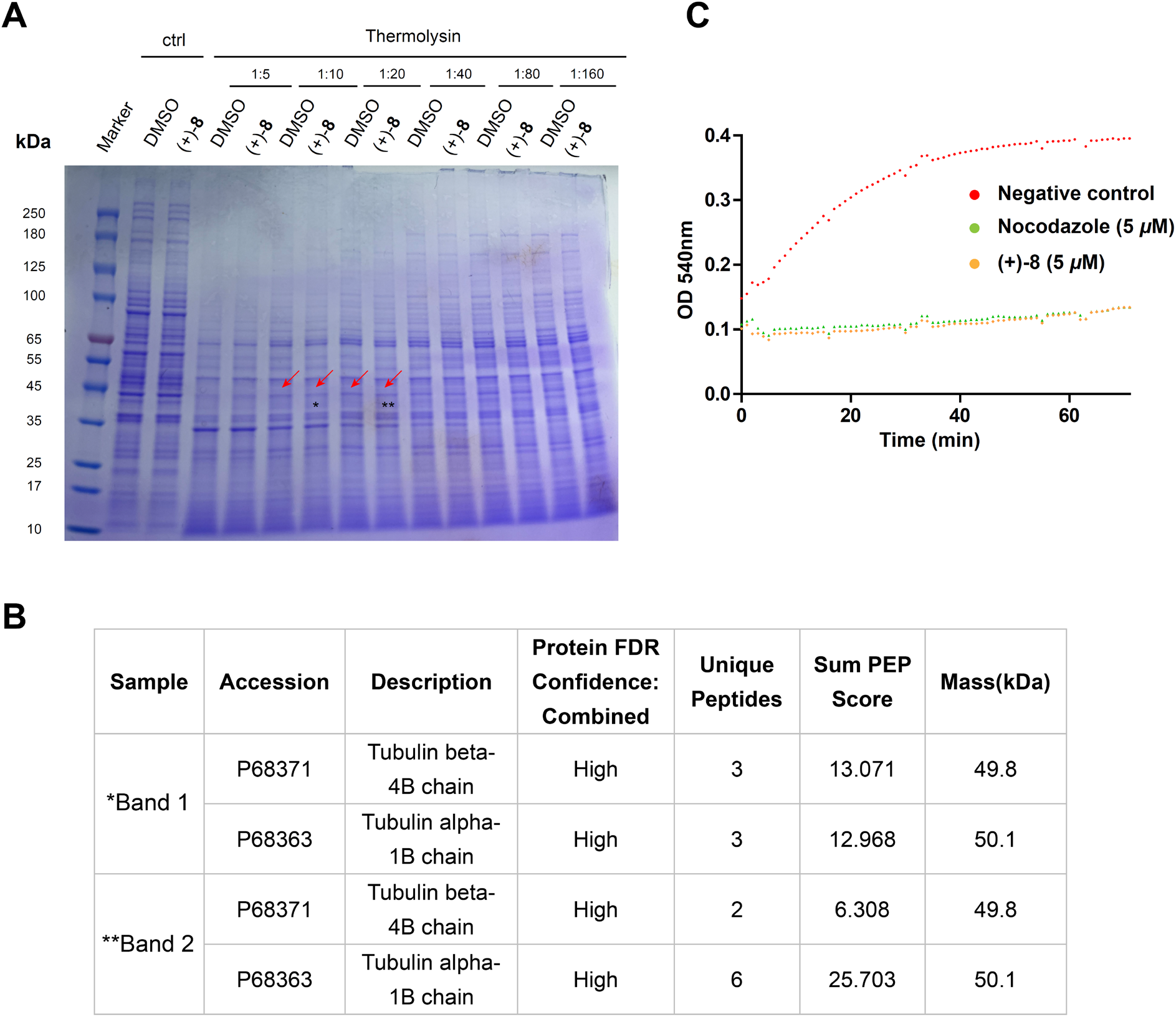
Identification of α- and β-tubulins as potential cellular targets of compound (+)-**8**. (A) A representative SDS-PAGE gel image showing the differential bands (ca. 45 kDa; marked with red arrows) in the DARTS assay to identify the target(s) of (+)-**8**. HeLa cell lysates were treated with (+)-**8** (10 *μ*M) for 1 h and then digested by thermolysin for 30 min. The asterisks indicate the bands subjected to mass spectrometry analysis. *Band 1. **Band 2. (B) Information on the potential protein targets of (+)-**8** identified by mass spectrometry. (C) Inhibition of in vitro tubulin polymerization by (+)-**8**. The curves for (+)-**8** (5.0 *μ*M), nocodazole (5.0 *μ*M), and the vehicle control (DMSO) are shown in yellow, green, and red, respectively.

### Compounds (+)-8 and (+)-9 target the colchicine-binding site of β-tubulin

We investigated the interaction between (+)-8 and tubulin in detail by X-ray crystallography analysis. This compound was soaked into a complex of two αβ-tubulin heterodimers, the stathmin-like domain (SLD) of RB3, and tubulin tyrosine ligase (TTL) (the T2R−TTL complex).[53] The structure of the protein−ligand complex was refined at 2.5 Å resolution (PDB ID: 7CPD; see Table S1 for data collection and refinement statistics).

As shown in Figure 5A, compound (+)-8 occupies the colchicine-binding site of β-tubulin (Chain B and Chain D) at the intradimer interface.[54–57] LigPlot-based protein−ligand interaction analysis[58] revealed the hydrophobic interaction of (+)-8 with residues such as β-Cys241, β-Leu242, β-Leu248, β-Ala250, β-Lys254, β-Leu255, β-Asn258, β-Met259, and β-Lys352 (Figure 5B). The indole N−H hydrogen atom forms a hydrogen bond with an oxygen atom of α-Thr179 (Figure 5B), highlighting the importance of the indole motif for binding to tubulin. The oxygen atom of the ketone forms a hydrogen bond with the N−H hydrogen atom of β-Asp251 (Figure 5B), which may explain the essential role of this oxygen atom observed in the SAR study. The interaction of the bromine atom with β-Asn350 (Figure 5B) could further enhance the binding of (+)-8 to β-tubulin,[59] consistent with the increased potency of this brominated analogue. The interaction pattern of (+)-9 in the binding pocket of Chain B (PDB ID: 7CPQ) is essentially identical to that of (+)-8, although the binding of the chlorinated analogue to Chain D was not constructed due to insufficient electron density (Figure S6). Comparison of the positions of MTAs in the colchicine-binding pocket revealed that (+)-8 and (+)-9 behave similarly to colchicine (PDB ID: 4O2B) but differently from nocodazole (PDB ID: 5CA1) (Figure 5C).

**Figure 5.**
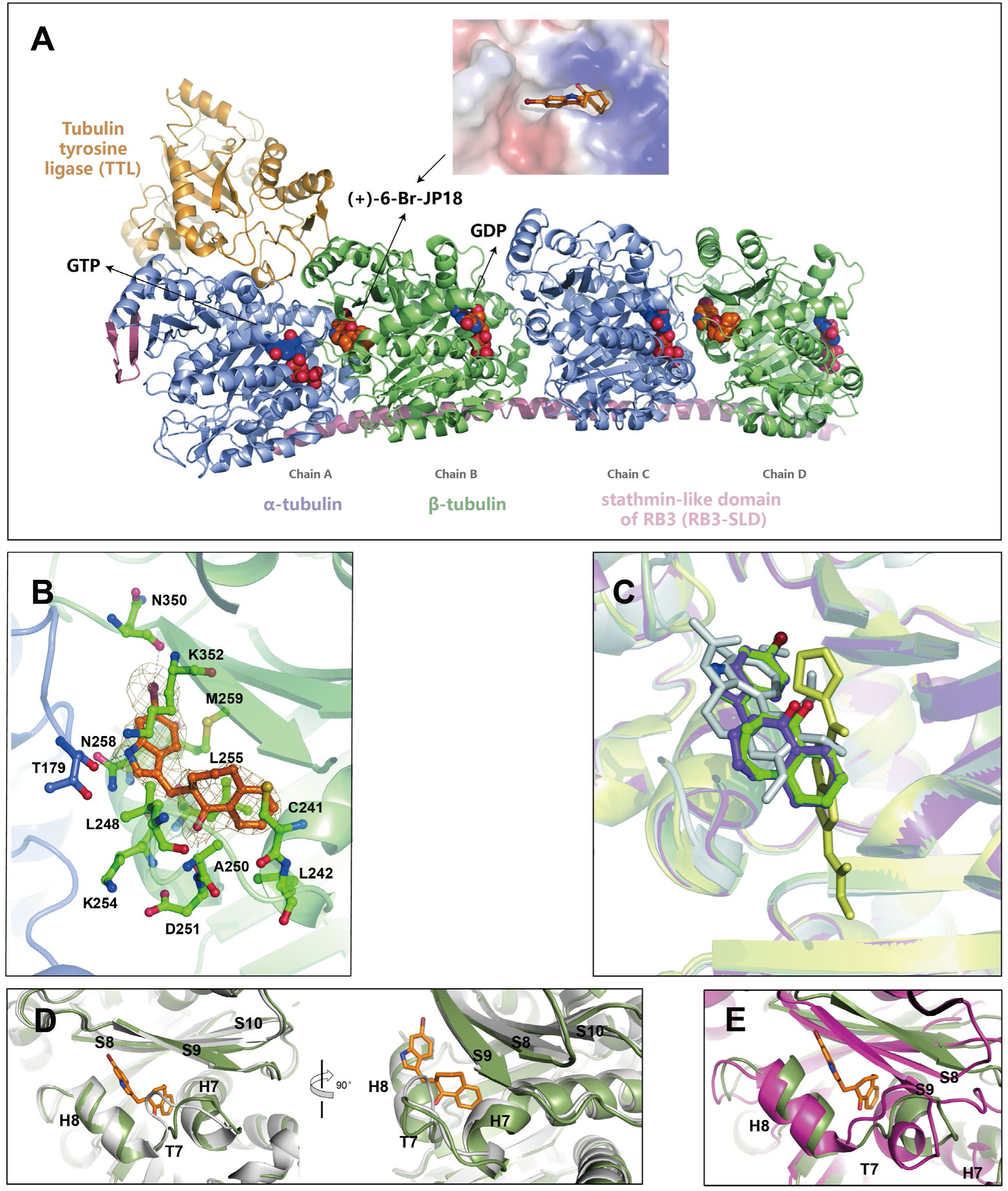
X-ray crystal structure of the T2R-TTL tubulin−(+)-**8** complex. (A) Overall structure of the T2R−TTL−(+)-**8** complex. α-Tubulin (light slate blue), β-tubulin (green), RB3-SLD (plum), and TTL (gold) are shown in cartoon representation. Compound (+)-**8**, GDP, and GTP are shown in sphere representation. (B) Close-up view of the interaction between (+)-**8** and the colchicine-binding site. α-Tubulin and β-tubulin are colored in light slate blue and green, respectively. The Fo−Fc simulated annealing omit map (contoured at 1.0 σ) is shown as a mesh. The residues involved in the interaction between (+)-**8** and β-tubulin are shown in stick representation. The hydrogen and halogen bonds are shown as dashed lines. (C) Comparison of the positions of (+)-**8** and (+)-**9** with those of colchicine and nocodazole in the colchicine-binding pocket. The four T2R-TTL-CBSI complex structures are superimposed, and (+)-**8** (green), (+)-**9** (medium purple), colchicine (gray), and nocodazole (yellow) are shown in stick representation. (D) Superimposition of the structure of (+)-**8**-bound β-tubulin (green) and that of unbound β-tubulin in the T2R−TTL complex (gray). (E) Superimposition of the structure of (+)-**8**-bound β-tubulin (green) and that of unbound β-tubulin in the tubulin sheets (violet-red).

We compared the structures of β-tubulin in the (+)-8-bound and unbound states. In the T2R−TTL complex (PDB ID: 4IIJ), where αβ-tubulin adopts the curved conformation, the side chains of Leu248 and Asn249 on the T7 loop of β-tubulin occlude the colchicine-binding site, as shown in Figure 5D. However, in the T2R−TTL−(+)-8 complex, the T7 loop of β-tubulin flips outward to accommodate ligand binding (Figure 5D). This conformational change is similar to that observed by Knossow and colleagues when colchicine binds.[60] Furthermore, superimposition of the structure of (+)-8-bound β-tubulin and that of β-tubulin in the tubulin sheets (PDB ID: 1JFF) (Figure 5E), where αβ-tubulin adopts the straight conformation, revealed that compound (+)-8 prevents the conversion from the curved conformation to the straight conformation by altering the conformations of the T7 loop, H7, H8, S8, and S9, thereby disrupting microtubule assembly. This is a characteristic mechanism of action of colchicine-binding site inhibitors (CBSIs).[61]

### Compounds (+)-8 and (+)-9 suppress the ubiquitin-mediated proteolysis of CENP-A

The uneven distribution of chromatids induced by (+)-8 and (+)-9 (Figure 3E) prompted us to investigate their effect on CENP-A, the deposition of which governs centromere positioning.[17–19] We observed that both compounds upregulated CENP-A in HeLa cells in a dose- and time-dependent manner [Figures 6A and 6F for (+)-8; Figures 6B and 6G for (+)-9]. To further elucidate the relationship between microtubule assembly and CENP-A regulation, we examined several other MTAs. Interestingly, not only the microtubule destabilizers nocodazole and vinblastine but also the microtubule stabilizer paclitaxel increased CENP-A levels in a dose- and time-dependent manner (Figures 6C–6E and Figures 6H–6J). These results suggested a significant correlation between microtubule dynamics and CENP-A regulation in mitosis.

**Figure 6.**
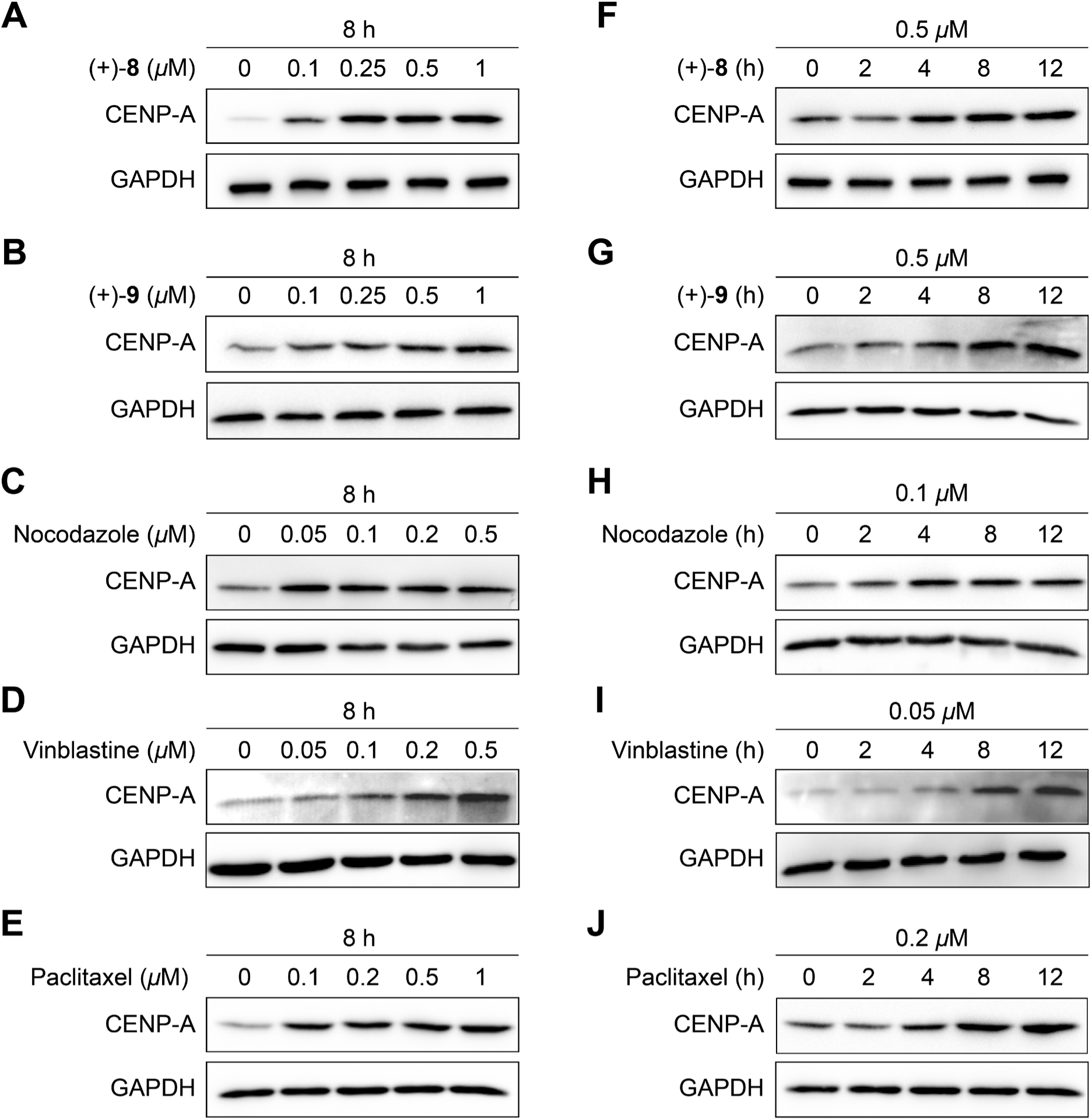
MTAs increased CENP-A levels. (A–E) Immunoblot analysis of CENP-A in HeLa cells treated with (+)-**8**, (+)-**9**, nocodazole, vinblastine, and paclitaxel, respectively, at indicated concentrations for 8h. (F–J) Immunoblot analysis of CENP-A in HeLa cells treated with (+)-**8** (0.50 *μ*M), (+)-**9** (0.50 *μ*M), nocodazole (0.10 *μ*M), vinblastine (0.050 *μ*M), and paclitaxel (0.10 *μ*M), respectively, for indicated durations. DMSO was used as a control for the compounds, and GAPDH was used as a loading control in immunoblotting.

We then explored whether MTAs affect CENP-A at the transcriptional or post-translational level. Quantitative real-time PCR analysis demonstrated that the tested MTAs did not influence the CENP-A mRNA levels in HeLa cells (Figure 7A). Therefore, we focused on the post-translational regulation of CENP-A, which may be mediated by the ubiquitin-proteasome system (UPS) and/or the lysosome system. The proteasome inhibitor MG132 increased CENP-A protein levels in HeLa cells (Figures 7B and 7C), while the lysosome inhibitors NH_4_Cl (Figure S7A) and chloroquine (CQ) (Figure S7B) had essentially no effect. Furthermore, overexpression of ubiquitin (Ub) downregulated CENP-A (Figures 7D and 7E), whereas overexpression of the ubiquitin-like proteins[62] neural precursor cell-expressed developmentally downregulated protein 8 (NEDD8) (Figure S7C) and small ubiquitin-like modifier 1 (SUMO1) (Figure S7D) did not. These observations indicated that ubiquitin-mediated proteolysis is the primary means of CENP-A degradation. Comparison of CENP-A levels in HeLa cells treated with the MTA or MG132 alone versus in those co-treated with the MTA and MG132 by immunoblotting showed that the combination did not further increase the CENP-A levels compared to (+)-8 or MG132 alone (Figures 7F and 7G). Similar observations were made with (+)-9 (Figures 7H and 7I) and nocodazole (Figure S7E). These results suggested that the tested MTAs act on the same CENP-A degradation system as the proteasome inhibitor.

**Figure 7.**
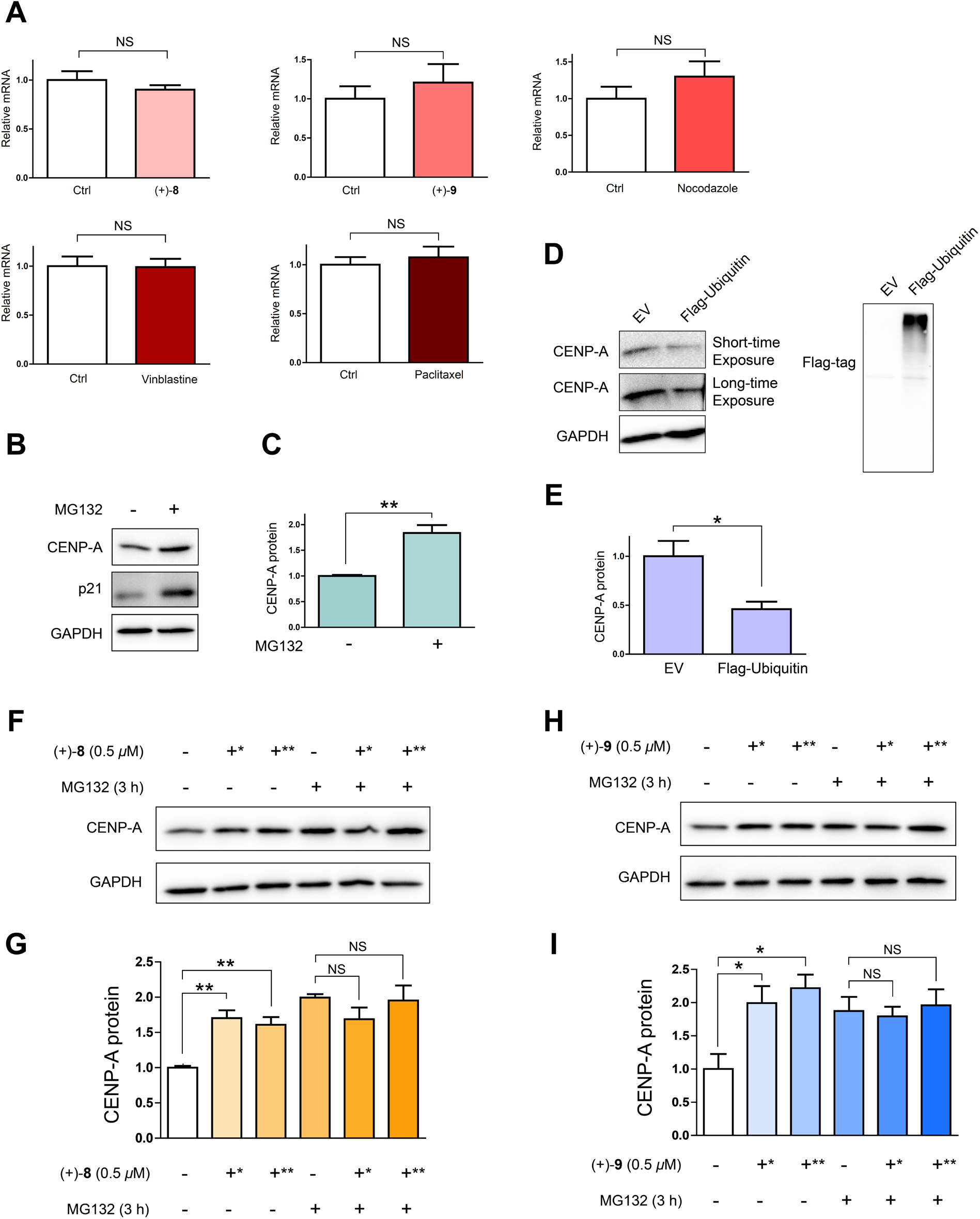
Compounds (+)-**8** and (+)-**9** suppress the degradation of CENP-A by inhibiting the ubiquitin-proteasome system. (A) qRT-PCR analysis of CENP-A mRNA levels in HeLa cells treated with (+)-**8** (0.50 *μ*M), (+)-**9** (0.50 *μ*M), nocodazole (0.10 *μ*M), vinblastine (0.050 *μ*M), and paclitaxel (0.10 *μ*M), respectively, for 6 h. β-actin mRNA levels were used for normalization. Data are representative of three independent biological replicates and are presented as mean ± s.e.m. NS = not significant (significance level: α = 0.05; *n* = 3, two-tailed Student’s *t*-test). (B) Immunoblot analysis of CENP-A in HeLa cells treated with MG132 (50 *μ*M) for 3 h. p21 served as a positive control. (C) Quantitative analysis of the immunoblotting data of CENP-A obtained from the above experiment. Data are representative of three independent biological replicates and are presented as mean ± s.e.m. \*\**P* < 0.01 (*n* = 3, two-tailed Student’s *t*-test). (D) Immunoblot analysis of CENP-A in HeLa cells transfected with FLAG-ubiquitin. (E) Quantitative analysis of the immunoblotting data of CENP-A obtained from the above experiment. Data are representative of three independent biological replicates and are presented as mean ± s.e.m. \**P* < 0.05 (*n* = 3, two-tailed Student’s *t*-test). (F) Comparison of CENP-A levels in HeLa cells treated with (+)-**8** (0.50 *μ*M) and co-treated with (+)-**8** (0.50 *μ*M) and MG-132 (50 *μ*M) for indicated durations by immunoblotting. *4 h. **6 h. (G) Quantitative analysis of the immunoblotting data of CENP-A obtained from the above experiment. Data are representative of three independent biological replicates and are presented as mean ± s.e.m. \*\**P* < 0.01, NS = not significant (*n* = 3, two-tailed Student’s *t*-test). (H) Comparison of CENP-A levels in HeLa cells treated with (+)-**9** (0.50 *μ*M) and co-treated with (+)-**9** (0.50 *μ*M) and MG-132 (50 *μ*M) for indicated durations by immunoblotting. *4 h. **6 h. (I) Quantitative analysis of the immunoblotting data of CENP-A obtained from the above experiment. Data are representative of three independent biological replicates and are presented as mean ± s.e.m. \**P* < 0.05, NS = not significant (*n* = 3, two-tailed Student’s *t*-test). DMSO was used as a control for the compounds, and GAPDH was used as a loading control in immunoblotting.

### The upregulation of CENP-A induced by (+)-8 is mediated by destabilizing the APC/C co-activator Cdh1

Anaphase-promoting complex/cyclosome (APC/C) is an important E3 ubiquitin ligase in the regulation of mitosis, whose substrate selectivity is determined by its co-activators Cdh1 and Cdc20.[63,64] Very recently, APC/C has been reported to participate in CENP-A regulation in *Drosophila*, where CENP-A^CID^ (the *Drosophila* orthologue of CENP-A) increased upon depletion of either Cdh1 or Cdc20.[65] In addition to APC/C, the cullin−RING E3 ubiquitin ligase (CRL) has also been implicated in CENP-A regulation. In HeLa cells, the CUL4−DDB1−DCAF11 E3 ubiquitin ligase mediates CENP-A degradation via poly-ubiquitylation,[27] whereas the CUL4A−RBX1−COPS8 E3 ubiquitin ligase directs CENP-A deposition via mono-ubiquitylation.[28] Building on these studies, we aim to identify the E3 ubiquitin ligase(s) involved in the MTA-induced upregulation of CENP-A.

We initially explored whether APC/C co-activators are associated with MTA-induced upregulation of CENP-A. Treatment with (+)-8 (Figures 8A and 8B) or paclitaxel (Figures 8C and 8D) resulted in a time-dependent decrease in Cdh1 levels and a time-dependent increase in CENP-A levels in HeLa cells. In contrast, Cdc20 remained unaffected upon such treatment (Figures 8A and 8C). Consistent observations in human hepatocellular carcinoma cells (Hep G2), lung carcinoma cells (A549), and normal liver cells (L-02) (Figure S8) supported that Cdh1 downregulation generally correlates with MTA-induced CENP-A upregulation. Comparison of Cdh1 degradation rates in HeLa cells treated with the protein synthesis inhibitor cycloheximide[66] (CHX) alone versus in those co-treated with (+)-8 and CHX revealed that compound (+)-8 downregulated Cdh1 at the post-translational level (Figure S9).

**Figure 8.**
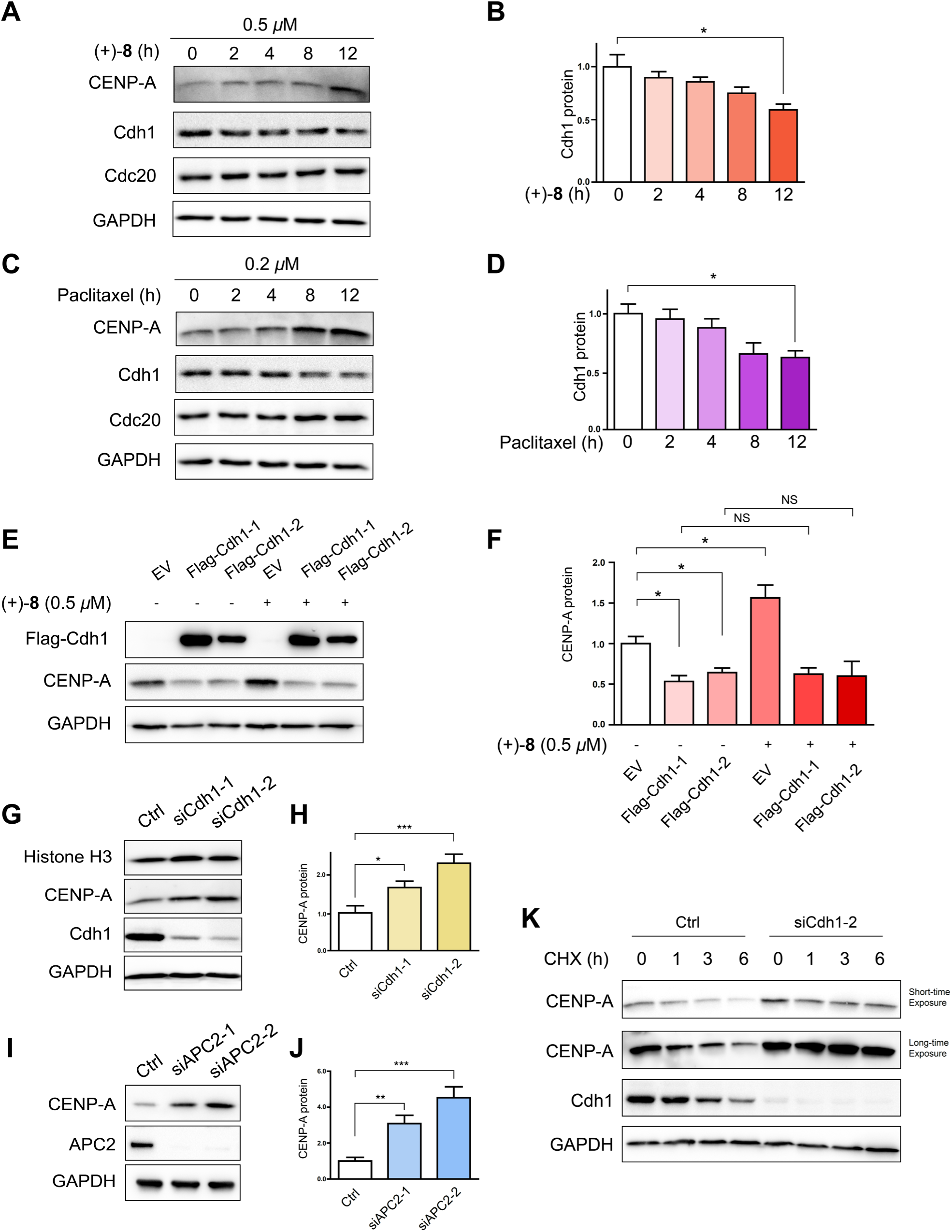
Compound (+)-**8** destabilizes Cdh1 to upregulate CENP-A. (A) Immunoblot analysis of CENP-A, Cdh1, and Cdc20 in HeLa cells treated with (+)-**8** (0.50 *μ*M) for indicated durations. (B) Quantitative analysis of the immunoblotting data of Cdh1 obtained from the above experiment. Data are representative of three independent biological replicates and are presented as mean ± s.e.m. \**P* < 0.05 (significance level: α = 0.05; *n* = 3, two-tailed Student’s *t*-test). (C) Immunoblot analysis of CENP-A, Cdh1, and Cdc20 in HeLa cells treated with paclitaxel (0.20 *μ*M) for indicated durations. (D) Quantitative analysis of the immunoblotting data of Cdh1 obtained from the above experiment. Data are representative of three independent biological replicates and are presented as mean ± s.e.m. \**P* < 0.05 (*n* = 3, two-tailed Student’s *t*-test). (E) Immunoblot analysis of CENP-A in HeLa cells transfected with FLAG-Cdh1 and then treated with (+)-**8** (0.50 *μ*M) for 12 h. Cdh1-1 and Cdh1-2 represent human Cdh1 isoform 1 (Q9UM11-2) and isoform 2 (Q9UM11-1), respectively. (F) Quantitative analysis of the immunoblotting data of CENP-A obtained from the above experiment. Data are representative of three independent biological replicates and are presented as mean ± s.e.m. \**P* < 0.05, NS = not significant (*n* = 3, two-tailed Student’s *t*-test). (G) Immunoblot analysis of CENP-A in HeLa cells transfected with siRNA targeting Cdh1 mRNA. (H) Quantitative analysis of the immunoblotting data of CENP-A obtained from the above experiment. Data are representative of three independent biological replicates and are presented as mean ± s.e.m. \**P* < 0.05, \*\*\**P* < 0.001 (*n* = 6, two-tailed Student’s *t*-test). (I) Immunoblot analysis of CENP-A in HeLa cells transfected with siRNA targeting APC2 mRNA. (J) Quantitative analysis of the immunoblotting data of CENP-A obtained from the above experiment. Data are representative of three independent biological replicates and are presented as mean ± s.e.m. \*\**P* < 0.01, \*\*\**P* < 0.001 (*n* = 6, two-tailed Student’s *t*-test). (K) Immunoblot analysis of CENP-A in HeLa cells transfected with siRNA targeting Cdh1 mRNA and then treated with CHX for indicated durations.

We then investigated the role of Cdh1 in CENP-A regulation. Overexpression of the Flag−Cdh1 construct in HeLa cells counteracted the effect of (+)-8 on the CENP-A level (Figures 8E and 8F). Furthermore, siRNA-mediated knockdown of Cdh1 increased CENP-A, without affecting its homologue histone H3 (Figures 8G and 8H). Conversely, Cdc20 knockdown had minimal effect on the CENP-A level (Figure S10). Consistent correlations between Cdh1 depletion and CENP-A upregulation observed across human hepatocellular carcinoma cells (Hep G2), lung carcinoma cells (A549), triple-negative breast cancer cells (MDA-MB-231), glioblastoma cells (U251), and normal liver cells (L-02) suggested that Cdh1 is generally responsible for CENP-A regulation in human cells (Figure S11). Comparison of CENP-A degradation rates in HeLa cells with and without Cdh1 knockdown (Figure 8K), using a CHX chase assay,[66] showed a significantly lower rate in the Cdh1-knockdown cells, which indicated that Cdh1 depletion upregulated CENP-A at the post-translational level. Additionally, the combination of Cdh1 knockdown and MTA treatment resulted in significant enhanced CENP-A accumulation compared to treatment with MTA alone (Figures S12A and S12B).

To confirm that Cdh1 functions as a co-activator of APC/C, we examined the effect of knocking down APC2, a core component of this E3 ubiquitin ligase, on CENP-A levels.[67] As shown in Figures 8I and 8J, CENP-A was upregulated in HeLa cells. Furthermore, the combination of APC2 depletion and treatment with (+)-8 considerably enhanced CENP-A accumulation compared to treatment with (+)-8 alone (Figure S13A). Paclitaxel exhibited a similar effect to (+)-8 (Figure S13B).

We also investigated the involvement of DCAF11, a key component of the CUL4-DDB1-DCAF11 E3 ubiquitin ligase, in MTA-induced CENP-A upregulation. Treatment with (+)-8 (Figure S14A) or paclitaxel (Figure S14B) did not significantly affect the levels of cullin 4A and DCAF11 levels in HeLa cells. Furthermore, the combination of DCAF11 knockdown and treatment with either MTA failed to further increase CENP-A levels compared to treatment with the MTA alone (Figures S12C and S12D). These results suggest that DCAF11 might not be involved in CENP-A degradation in this context.

## • **▪** DISCUSSION

In this study, we identified β-tubulin as a cellular target of the cell cycle inhibitors (+)-6-Br-JP18 and (+)-6-Cl-JP18 by using the DARTS-based method and revealed the interaction between these small molecules and the colchicine-binding site of β-tubulin through X-ray crystallography analysis (Figure 9). Despite being an older class of anticancer agents, CBSIs have regained interest in anticancer drug discovery in recent years. This resurgence is presumably due to the promising progress of combretastatin analogues such as fosbretabulin (CA-4P) and OXi-4503 in clinical trials[54,55] and the successful use of MTAs as payloads of antibody−drug conjugates.[68] However, combretastatins suffer from configurational instability due to *cis*/*trans* isomerization, while other promising CBSIs in clinical trials, such as plinabulin and lisavanbulin, pose a challenge for further chemical modifications because of their complex structures. The 6-halo-JP18 scaffold has the potential to address these obstacles and could be valuable in therapeutic development.

**Figure 9.**
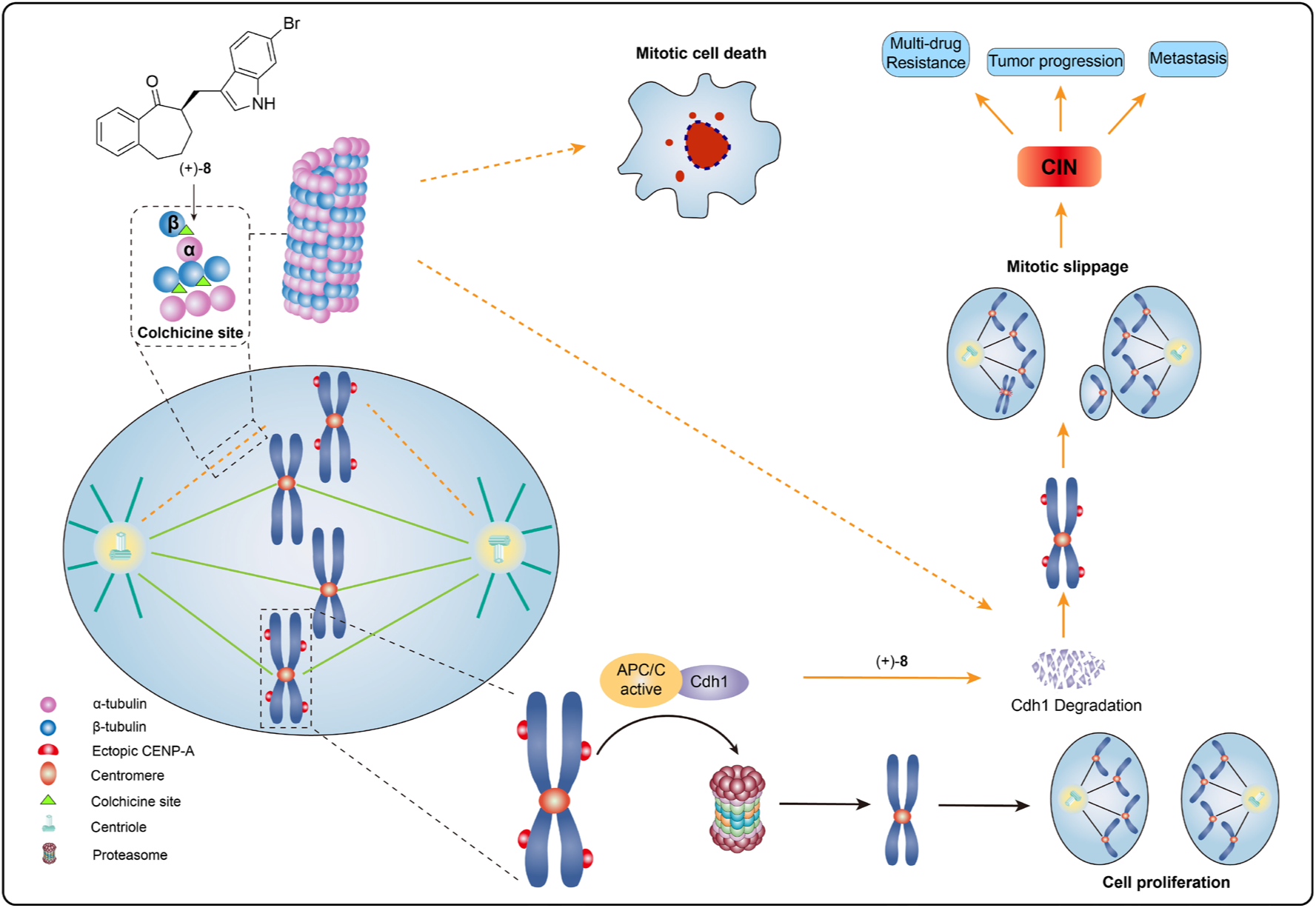
Schematic illustrating the mechanism of action of (+)-6-Br-JP18 on spindle microtubule assembly and Cdh1-mediated CENP-A proteolysis. (+)-6-Br-JP18 disrupts microtubule assembly by targeting the colchicine-binding site of β-tubulin, which leads to downregulation of Cdh1, a co-activator of the APC/C E3 ubiquitin ligase, and accumulation of its substrate CENP-A. This accumulation impairs the accuracy of chromosome segregation in mitosis and may result in mitotic slippage and chromosome instability.

We elucidated the crucial role of Cdh1 in CENP-A regulation in human cells (Figure 9). Maintaining centromeric identity necessitates precise deposition of CENP-A.[14–16] Aberrant overexpression of CENP-A, which leads to its mislocalization, is associated with the malignant potential of tumors.[21–24] Therefore, understanding the regulatory mechanisms of CENP-A expression and degradation could shed light on cancer development. Ubiquitin-mediated proteolysis of CENP-A regulates its stability and prevents ectopic assembly of CENP-A chromatin across species.[25,26] In *Drosophila*, the E3 ubiquitin ligases APC/C^Cdh1^ and SCF^Ppa^ target CENP-A^CID^ in G1 phase and G1/S phases, respectively.[65] We discovered that Cdh1 downregulates CENP-A in a “necessary and sufficient” manner during mitosis in human cells. This represents an advancement in understanding the regulatory mechanism of CENP-A in mammalian cells.

We established a link between microtubule dynamics and Cdh1-mediated CENP-A proteolysis using MTAs, including (+)-6-Br-JP18 and (+)-6-Cl-JP18, as tools (Figure 9). Despite being an effective class of chemotherapeutic agents, MTAs can induce drug resistance and disease relapse after long-term use.[7,69] An important mechanism contributing to this resistance is cancer cells evading drug-induced mitotic arrest to avoid subsequent cell death.[7,8] This process, also known as mitotic slippage, is closely associated with CIN,[8] which can result from aberrant overexpression of CENP-A.[19,20] Our discovery that disrupting spindle microtubule assembly with MTAs leads to CENP-A accumulation fills a gap in the possible mechanism for the MTA-induced drug resistance and suggests a potential role for Cdh1 in preventing the emergence of such resistance in cancer therapy.

## ACKNOWLEDGMENTS

This paper is dedicated to the memory of Prof. Hualiang Jiang. We thank Profs. Yang Wang, Kuiling Ding, Xin Zhang, Linghua Meng, and Mr. Fahui Zhu for helpful discussions and Institutes of Biomedical Science (IBS) at Fudan University and Shanghai Synchrotron Radiation Facility (SSRF) for technical assistance. This work was supported by National Natural Science Foundation of China (21931014, 21572064, and U2002221), Chinese Academy of Sciences (YSBR-095, XBZG-ZDSYS-202303, 121731KYSB20190039, and QYZDB-SSW-SLH040), and Science and Technology Commission of Shanghai Municipality (JCYJ-SHFY-2022-005 and 20430713400). A.L. is grateful to the New Cornerstone Science Foundation for the XplorerPrize. The original results described in this paper were uploaded to the preprint server for biology, bioRxiv (https://doi.org/10.1101/2023.04.25.538200), on April 26, 2023.

## REFERENCES

[1] T. J. Mitchison, E. D. Salmon, Nat. Cell Biol. 2001, 3, E17–E21.

[2] J. R. McIntosh, Cold Spring Harb. Perspect. Biol. 2016, 8, a023218.

[3] D. A. Compton, Annu. Rev. Biochem. 2000, 69, 95–114.

[4] N. B. Gudimchuk, J. R. McIntosh, Nat. Rev. Mol. Cell Biol. 2021, 22, 777–795.

[5] M. A. Jordan, L. Wilson, Nat. Rev. Cancer 2004, 4, 253–265.

[6] V. Cermak, V. Dostal, M. Jelinek, L. Libusova, J. Kovar, D. Rosel, J. Brabek, Microtubule-targeting agents and their impact on cancer treatment. Eur. J. Cell Biol. 2020, 99, 151075.

[7] B. Cheng, K. Crasta, Endocr. Relat. Cancer 2017, 24, T97–T106.

[8] D. Sinha, P. H. G. Duijf, K. K. Khanna, Cell Cycle 2019, 18, 7–15.

[9] S. E. McClelland, R. A. Burrell, C. Swanton, Cell Cycle 2009, 8, 3262–3266.

[10] R. Vishwakarma, K. J. McManus, Cancers 2020, 12, 824.

[11] D. A. Lukow, J. M. Sheltzer, Trends Cancer 2022, 8, 43–53.

[12] M. R. Ippolito, V. Martis, S. Martin, A. E. Tijhuis, C. Hong, R. Wardenaar, M. Dumont, J. Zerbib, D. C. J. Spierings, D. Fachinetti, U. Ben-David, F. Foijer, S. Santaguida, Dev. Cell 2021, 56, 2440–2454.

[13] J. M. Reploglea, W. Zhou, A. E. Amaroa, J. M. McFarlandc, M. Villalobos-Ortizd, J. Ryan, A. Letai, O. Yilmaz, J. Sheltzer, S. J. Lippard, U. Ben-David, A. Amon, Proc. Natl. Acad. Sci. USA 2020, 117, 30566–30576.

[14] B. E. Black, E. A. Bassett, Curr. Opin. Cell Biol. 2008, 20, 91–100.

[15] V. De Rop, A. Padeganeh, P. S. Maddox, Chromosoma 2012, 121, 527–538.

[16] K. Kixmoeller, P. K. Allu, B. E. Black, Open Biol. 2020, 10, 200051.

[17] C. C. Chen, B. G. Mellone, J. Cell Biol. 2016, 214, 13–24.

[18] K. L. McKinley, I. M. Cheeseman, Nat. Rev. Mol. Cell Biol. 2016, 17, 16–29.

[19] M. A. Mahlke, Y. Nechemia-Arbely, Genes 2020, 11, 810.

[20] R. L. Shrestha, G. S. Ahn, M. I. Staples, K. M. Sathyan, T. S. Karpova, D. R. Foltz, M. A. Basrai, Oncotarget 2017, 8, 46781–46800.

[21] L. M. Zink, S. B. Hake, Curr. Opin. Genet. Dev. 2016, 37, 82–89.

[22] A. B. Sharma, S. Dimitrov, A. Hamiche, E. Van Dyck, Nucleic Acids Res. 2019, 47, 1051–1069.

[23] S. L. McGovern, Y. Qi, L. Pusztai, W. F. Symmans, T. A. Buchholz, Breast Cancer Res. 2012, 14, R72.

[24] X. Sun, P.-L. Clermont, W. Jiao, C. D. Helgason, P. W. Gout, Y. Wang, S. Qu, Int. J. Cancer 2016, 139, 899–907.

[25] Y. Niikura, R. Kitagawa, K. Kitagawa, Dev. Cell 2017, 40, 7–8.

[26] S. Srivastava, D. R. Foltz, Chromosoma 2018, 127, 279–290.

[27] K. Wang, Y. Liu, Z. Yu, B. Gu, J. Hu, L. Huang, X. Ge, L. Xu, M. Zhang, J. Zhao, M. Hu, R. Le, Q. Wu, S. Ye, S. Gao, X. Zhang, R. M. Xu, G. Li, Cell Rep. 2021, 37, 109987.

[28] Y. Niikura, R. Kitagawa, H. Ogi, R. Abdulle, V. Pagala, K. Kitagawa, Dev. Cell 2015, 32, 589–603.

[29] I. S. Marcos, R. F. Moro, I. Costales, P. Basabe, D. Diez, Nat. Prod. Rep. 2013, 30, 1509–1526.

[30] M. A. Corsello, J. Kim, N. K. Garg, Chem. Sci. 2017, 8, 5836–5844.

[31] N. Devi, K. Kaur, A. Biharee, V. Jaitak, Anticancer Agents Med. Chem. 2021, 21, 1802–1824.

[32] T. V. Sravanthi, S. L. Manju, Eur. J. Pharm. Sci. 2016, 91, 1–10.

[33] A. A. Sallam, N. M. Ayoub, A. I. Foudah, C. R. Gissendanner, S. A. Meyer, K. A. El Sayed, *Eur.* J. Med. Chem. 2013, 70, 594–606.

[34] A. A. Sallam, W. E. Houssen, C. R. Gissendanner, K. Y. Orabi, A. I. Foudah, K. A. El Sayed, MedChemComm 2013, 4, 1360–1369.

[35] A. A. Goda, A. B. Siddique, M. Mohyeldin, N. M. Ayoub, K. A. El Sayed, Mar. Drugs. 2018, 16, 157.

[36] Y. Sun, R. Li, W. Zhang, A. Li, Angew. Chem. Int. Ed. 2013, 52, 9201–9204.

[37] Y. Sun, P. Chen, D. Zhang, M. Baunach, C. Hertweck, A. Li, Angew. Chem. Int. Ed. 2014, 53, 9012– 9016.

[38] Z. Lu, M. Yang, P. Chen, X. Xiong, A. Li, Angew. Chem. Int. Ed. 2014, 53, 13840–13844.

[39] Z. Meng, H. Yu, L. Li, W. Tao, H. Chen, M. Wan, P. Yang, D. J. Edmonds, J. Zhong, A. Li, Nat. Commun. 2015, 6, 6096.

[40] S. Zhou, H. Chen, Y. Luo, W. Zhang, A. Li, Angew. Chem. Int. Ed. 2015, 54, 6878–6882.

[41] Z. Lu, H. Li, M. Bian, A. Li, J. Am. Chem. Soc. 2015, 137, 13764–13767.

[42] X. Xiong, D. Zhang, J. Li, Y. Sun, S. Zhou, M. Yang, H. Shao, A. Li, Chem. Asian J. 2015, 10, 869– 872.

[43] H. Li, Q. Chen, Z. Lu, A. Li, J. Am. Chem. Soc. 2016, 138, 15555–15558.

[44] J. Pei, S. Zhou, F. Yang, Y. Sun, A. Li, W. D. Zhang, W. He, Chem. Asian J. 2016, 11, 2715–2718.

[45] J. Kamenz, S. Hauf, Trends Cell Biol. 2017, 27, 42–54.

[46] Y.-P. Li, Z.-Q. Li, B. Zhou, M.-L. Li, X.-S. Xue, S.-F. Zhu, Q.-L. Zhou, ACS Catal. 2019, 9, 6522– 6529.

[47] S. J. Nowak, V. G. Corces, Trends Genet. 2004, 20, 214–220.

[48] C. F. A. Warren, M. W. Wong-Brown, N. A. Bowden, Cell Death Dis. 2019, 10, 177.

[49] E. P. Rogakou, D. R. Pilch, A. H. Orr, V. S. Ivanova, W. M. Bonner, J. Biol. Chem. 1998, 273, 5858– 5868.

[50] S. H. Kaufmann, S. Desnoyers, Y. Ottaviano, N. E. Davidson, G. G. Poirier, Cancer Res. 1993, 53, 3976–3985.

[51] B. Lomenick, R. Hao, N. Jonai, R. M. Chin, M. Aghajan, S. Warburton, J. Wang, R. P. Wu, F. Gomez, J. A. Loo, J. A. Wohlschlegel, T. M. Vondriska, J. Pelletier, H. R. Herschman, J. Clardy, C. F. Clarke, J. Huang, Proc. Natl. Acad. Sci. USA 2009, 106, 21984–21989.

[52] M. Y. Pai, B. Lomenick, H. Hwang, R. Schiestl, W. McBride, J. A. Loo, J. Huang, Methods Mol. Biol. 2015, 1263, 287–298.

[53] A. E. Prota, K. Bargsten, D. Zurwerra, J. J. Field, J. F. Diaz, K. H. Altmann, M. O. Steinmetz, Science 2013, 339, 587–590.

[54] E. C. McLoughlin, N. M. O’Boyle, Pharmaceuticals 2020, 13, 8.

[55] Y. Duan, W. Liu, L. Tian, Y. Mao, C. Song, Curr. Top. Med. Chem. 2019, 19, 1289–1304.

[56] P. Zhou, Y. Liu, L. Zhou, K. Zhu, K. Feng, H. Zhang, Y. Liang, H. Jiang, C. Luo, M. Liu, Y. Wang, J. Med. Chem. 2016, 59, 10329–10334.

[57] P. Zhou, Y. Liang, H. Zhang, H. Jiang, K. Feng, P. Xu, J. Wang, X. Wang, K. Ding, C. Luo, M. Liu, Y. Wang, Eur. J. Med. Chem. 2018, 144, 817–842.

[58] A. C. Wallace, R. A. Laskowski, J. M. Thornton, Protein Eng. 1995, 8, 127–134.

[59] Y. Lu, Y. Liu, Z. Xu, H. Li, H. Liu, W. Zhu, Expert Opin. Drug Discov. 2012, 7, 375–383.

[60] R. B. G. Ravelli, B. Gigant, P. A. Curmi, I. Jourdain, S. Lachkar, A. Sobel, M. Knossow, Nature 2004, 428, 198–202.

[61] J. Wang, D. D. Miller, W. Li, Drug Discov. Today 2022, 27, 759–776.

[62] A. G. van der Veen, H. L. Ploegh, Annu. Rev. Biochem. 2012, 81, 323–357.

[63] R. van Leuken, L. Clijsters, R. Wolthuis, Biochim. Biophys. Acta 2008, 1786, 49–59.

[64] M. S. Schrock, B. R. Stromberg, L. Scarberry, M. K. Summers, Semin. Cancer Biol. 2020, 67, 80– 91.

[65] O. Moreno-Moreno, M. Torras-Llort, F. Azorin, Nucleic Acids Res. 2019, 47, 3395–3406.

[66] P. Zhou, Methods Mol. Biol. 2004, 284, 67–77.

[67] L. K. Teixeira, S. I. Reed, Annu. Rev. Biochem. 2013, 82, 387–414.

[68] Z. Wang, H. Li, L. Gou, W. Li, Y. Wang, Acta Pharm. Sin. B 2023, 13, 4025–4059.

[69] L. Mosca, A. Ilari, F. Fazi, Y. G. Assaraf, G. Colotti, Drug Resist. Updat. 2021, 54, 100742.

