## Supplemental file for "Target Identification and Mechanistic Characterization of Indole Terpenoid Mimics: Proper Spindle Microtubule Assembly Is Essential for Cdh1-Mediated Proteolysis of CENP-A"

Electronic Supporting Information

**I	Materials and Methods**

**II	Figures and Tables**

**III	References**

**I	Materials and Methods**

**Preparation of JP18 analogues.** JP18 analogues were prepared in racemic form through a previously developed conjugate addition approach.^1–3^

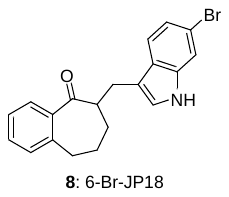

**6-Br-JP18 (8):** This compound was obtained as a white foam. **8**: *R*_f_ = 0.23 (silica gel, EtOAc:petroleum ether 1:7); IR (film): ν_max_ = 3422, 2933, 2859, 1734, 1672, 1597, 1453, 1334, 1276, 1254, 1048, 802, 737 cm^−1^; ^1^H NMR (400 MHz, CDCl_3_): δ = 7.99 (br s, 1 H), 7.62 (d, *J* = 7.5 Hz, 1H), 7.51–7.42 (m, 2 H), 7.40–7.32 (m, 1 H), 7.25 (t, *J* = 7.4 Hz, 1 H), 7.22–7.13 (m, 2 H), 6.95 (s, 1 H), 3.37 (dd, *J* = 14.5, 6.4 Hz, 1 H), 3.29–3.14 (m, 1 H), 3.04–2.84 (m, 3 H), 2.10–1.91 (m, 2 H), 1.76–1.54 (m, 2 H) ppm; ^13^C NMR (101 MHz, CDCl_3_): δ = 207.4, 142.2, 140.0, 137.0, 131.4, 129.9, 128.2, 126.5, 126.4, 123.0, 122.5, 120.1, 115.4, 114.5, 114.0, 50.7, 33.6, 30.3, 26.3, 25.5 ppm; HRMS (*m/z*): calcd for C_20_H_18_BrNONa^+^ ([M + Na]^+^): 390.0464, found: 390.0459.

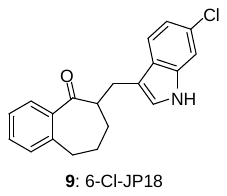

**6-Cl-JP18 (9):** This compound was obtained as a white foam. **9**: *R*_f_ = 0.33 (silica gel, EtOAc:petroleum ether 1:4); IR (film): ν_max_ = 3424, 2935, 2859, 1678, 1619, 1597, 1455, 1336, 804, 737 cm^−1^; ^1^H NMR (400 MHz, CDCl_3_): δ = 7.96 (br s, 1 H), 7.61 (d, *J* = 7.7 Hz, 1 H), 7.49 (d, *J* = 8.5 Hz, 1 H), 7.40–7.32 (m, 1 H), 7.32 (d, *J* = 1.6 Hz, 1 H), 7.28–7.22 (m, 1 H), 7.19 (d, *J* = 7.5 Hz, 1 H), 7.06 (dd, *J* = 8.5, 1.7 Hz, 1 H), 6.97 (d, *J* = 1.9 Hz, 1 H), 3.38 (dd, *J* = 14.6, 6.4 Hz, 1 H), 3.26–3.15 (m, 1 H), 3.04–2.89 (m, 2 H), 2.90 (dd, *J* = 14.6, 7.2 Hz, 1 H), 2.09–1.92 (m, 2 H), 1.75–1.59 (m, 2 H) ppm; ^13^C NMR (101 MHz, CDCl_3_): δ = 207.4, 142.2, 140.1, 136.6, 131.4, 130.0, 128.3, 128.0, 126.5, 126.4, 123.1, 120.1, 119.9, 114.7, 111.1, 50.8, 33.7, 30.4, 26.4, 25.6 ppm; HRMS (*m/z*): calcd for C_20_H_18_ClNONa^+^ ([M + Na]^+^): 346.0969, found: 346.0966.

**Antibodies.**

| Antibody | Vendor | Catalog No. |
| --- | --- | --- |
| Rabbit anti-Cyclin B1 (D5C10) Antibody | ABclonal | A19037 |
| Mouse anti-Cdc2 (POH1) Antibody | CST | 9116 |
| Rabbit anti-Phospho-Cdc2 (Tyr15) Antibody | CST | 4539 |
| Rabbit anti-Phospho-Histone H3 (S10) Antibody | CST | 3377 |
| Rabbit anti-Phospho-Histone H3 (S28) Antibody | ABclonal | AP0839 |
| Rabbit anti-Histone H3 Antibody | ABclonal | A2348 |
| Rabbit anti-Phospho-Histone H2A.X (Ser139) Antibody | Abcam | ab81299 |
| Rabbit anti-Bcl-2 Antibody | Abways | CY5032 |
| Rabbit anti-Cleaved PARP Antibody | Bimake | A5034 |
| Mouse anti-α-Tubulin Antibody | Sigma−Aldrich | T9026 |
| Rabbit anti-CENP-A Antibody | CST | 2186 |
| Rabbit anti-CENP-A Antibody | Abcam | Ab45694 |
| Rabbit anti-GAPDH Antibody | Abways | AB0037 |
| Rabbit anti-CDKN1A/P21CIP1 Antibody | ABclonal | A21772 |
| Rabbit anti-LC3B Antibody | Sigma−Aldrich | L7543 |
| Mouse anti-FLAG-Tag Antibody | Affinity | T0003 |
| Mouse anti-FLAG-Tag Antibody | Bimake | A5712 |
| Rabbit anti-Cdc20 Antibody | ABclonal | A15656 |
| Mouse anti-Cdh1 (Ab-2) Antibody | Sigma−Aldrich | CC43 |
| Mouse anti-Myc-Tag (9B11) Antibody | CST | 2276 |
| Rabbit anti-Cullin 4A Antibody | CST | 2699 |
| Rabbit anti-DCAF11 Antibody | ABclonal | A15519 |
| Rabbit anti-APC2 Antibody | Proteintech | 13559-1-AP |
| Peroxidase-AffiniPure Goat anti-Rabbit IgG (H+L) | Jackson ImmunoReserch | 147832 |
| Peroxidase-AffiniPure Goat anti-Mouse IgG (H+L) | Jackson ImmunoReserch | 139283 |
| Goat anti-Rabbit IgG (H+L) (Alexa Fluor 647) | Abcam | ab150079 |
| Goat anti-Mouse IgG (H+L) (Alexa Fluor 488) | Abcam | ab150113 |

**Reagents.** MG-132 was purchased from Selleck Chemicals. Nocodazole, vinblastine, paclitaxel, chloroquine (CQ), and cycloheximide (CHX) were purchased from MedChemExpress. NH_4_Cl was purchased from Adamas-beta.

**Cell culture.** Human cell lines HeLa, MDA-MB-231, A549, U251, Hep G2, and L-02 cells were purchased from the Cell Bank of Chinese Academy of Sciences. HeLa, Hep G2, and U251 cells were cultured in HyClone Dulbecco’s Modified Eagle’s Medium (DMEM) with high glucose (Cytiva) supplemented with 10% fetal bovine serum (FBS; Biological Industries) and 1% penicillin−streptomycin (Gibco, Thermos Fisher Scientific). MDA-MB-231, A549, and L-02 cells were cultured in HyClone RPMI 1640 Medium (Cytiva) supplemented with 10% FBS and 1% penicillin−streptomycin. Cells were maintained at 37 °C with 5% CO_2_ in a humidified incubator.

**Plasmids.** Human ubiquitin, SUMO1, and NEDD8 were cloned into mammalian expression vector pcDNA3.1(+) (Thermo Fisher Scientific), respectively. The FLAG sequence was inserted into the N-terminus of ubiquitin. The Myc sequence was inserted into the N-termini of SUMO1 and NEDD8. Human Cdh1 isoform 1 (UniProt ID: Q9UM11-2) and isoform 2 (UniProt ID: Q9UM11-1) were cloned into mammalian expression vector pCMV-N-FLAG (Addgene), respectively.

**Transfection.** Transfection was performed by using Lipofectamine 2000 (Thermo Fisher Scientific) for plasmids and Lipofectamine 3000 (Thermo Fisher Scientific) for siRNA. Cells were subjected to analysis 36–48 h after transfection. siRNAs were purchased from GenePharma.

| siRNA | Sequence |
| --- | --- |
| non-relevant siRNA | UUCUCCGAACGUGUCACGUTT |
| siRNA targeting Cdc20 | No. 1: AACGGCAGGACUCCGGGCCGATT  No. 2: AAUGGCCAGUGGUGGUAAUGATT |
| siRNA targeting Cdh1 | No. 1: GCCAGATCGTCATCCAGAA  No. 2: CCAACTGGAGCGTGAACTT |
| siRNA targeting APC2 | No. 1: ACAUAGUGUGGACUUCUUCUC  No. 2: AUAUAGACUCUCAAGAAGCAC |
| siRNA targeting DCAF11 | No. 1: CCGGGAGCUGGAAUUCAAU  No. 2: GCUACUCUCAGAAGGCUUU |

**Cell viability assay.** HeLa cells were seeded in a 96-well plate at a density of 1.0 × 10^4^ per well and treated with the tested compound at concentrations ranging from 0 to 10 *μ*M for 24 h. Dimethyl sulfoxide (DMSO; Sigma−Aldrich) was used as the vehicle for the compound. Cell viability was analyzed by using the Cell Counting Kit-8 (CCK-8; Dojindo Molecular Technologies).

**Cell cycle analysis.** HeLa cells were seeded in a 6-well plate at a density of 1.0 × 10^6^ per well and treated with the tested compound at the indicated concentrations for 8 h. DMSO was used as the vehicle for the compound. The cells were stained by using the Cell Cycle and Apoptosis Analysis Kit (Beyotime). Cellular DNA content was measured with a CytoFLEX S flow cytometer (Beckman Coulter), and the data obtained were analyzed with FlowJo 7.6.

**Immunoblot analysis.** Cells were lysed in RIPA lysis buffer (Cell Signaling Technology) supplemented with Protease Inhibitor Cocktail (Roche), Phosphatase Inhibitor Cocktail (Bimake), and phenylmethanesulfonyl fluoride (PMSF; Beyotime). Proteins from the cell lysate were separated by sodium dodecyl sulfate-polyacrylamide gel electrophoresis (SDS-PAGE) and transferred to a polyvinylidene difluoride (PVDF) membrane (Sigma−Aldrich). The PVDF membrane was blocked with 5% skim milk in Tris-buffered saline containing 0.1% Tween 20 (TBST) and incubated with the corresponding primary and secondary antibodies. The membrane was then incubated with Tanon High-sig ECL Western Blotting Substrate (Tanon), and the immunoreactive bands on the membrane were analyzed by using Tanon 5200 Chemiluminescent Imaging System (Tanon). Quantitative analysis of the immunoblot data was performed with Image J.

**Immunofluorescent analysis.** HeLa cells cultured on glass-bottom culture dishes (NEST) were fixed with 4% paraformaldehyde for 10 min and permeabilized with phosphate buffered saline (PBS) containing 10% FBS and 0.5% saponin at room temperature for 10 min. Samples were incubated with the primary and secondary antibodies shown above and mounted by using Fluoroshield with DAPI (Sigma−Aldrich). Fluorescence images were captured with a Leica Stellaris 5 SR laser confocal microscope and analyzed with Leica Application Suite X (LAS X).

**Drug affinity responsive target stability (DARTS)-based target identification.** Huang’s protocol^4,5^ was used for this experiment. HeLa cells were lysed with M-PER lysis buffer (Thermo Fisher Scientific). The cell lysate was divided into identical aliquots and incubated with compound (+)-**8** at various concentrations or vehicle control (DMSO) at room temperature for 1 h. The samples were then incubated with thermolysin at various dilutions (1:5–1:160) at room temperature for 30 min before they were separated by SDS-PAGE and stained with Coomassie Blue. The protein bands were analyzed by mass spectrometry at Institute of Biomedical Science, Fudan University.

**Tubulin polymerization assay.** The tubulin polymerization assay was performed by using the HTS-Tubulin Polymerization Assay Biochem Kit (Cytoskeleton).

**Protein expression and purification.** The stathmin-like domain of RB3 (RB3-SLD) and tubulin tyrosine ligase (TTL) were obtained as reported.^6^ RB3-SLD was cloned into the pET22b vector and overexpressed in *Escherichia* coli. The protein was purified by anion-exchange chromatography on a Q Sepharose Fast Flow column (GE Healthcare) and fast protein liquid chromatography on a Superose 12 HR 10/30 gel filtration column (Amersham Pharmacia Biotech) and then concentrated to 10 mg/mL. TTL was cloned into the pET22b vector with a hexahistidine tag. The protein was purified by immobilized metal affinity chromatography on a HisTrap HP column (GE Healthcare) and size exclusion chromatography on a Superdex 200 column (GE Healthcare) and then concentrated to 10 mg/mL. Bovine brain tubulin was purchased from Cytoskeleton.

**Crystallization and crystal soaking.** The T2R−TTL complex was formed by mixing tubulin, RB3-SLD, and TTL in a ratio of 2:1.3:1.2.^6^ The complex was concentrated to 20 mg/mL before the addition of 1.0 mM AMPPCP, 5.0 mM tyrosinol, and 10 mM DTT and crystallized by sitting-drop vapor diffusion at 16 ℃. The crystals grew in a series of well solutions consisting of 6–10% PEG 4000, 4–10% glycerol, 30 mM MgCl_2_, and 100 mM MES/imidazole at pH 6.5–6.9 and reached their maximum dimensions within 4 d. The resulting crystals were soaked in a well solution containing the tested compound at 1.0 mM for 12–24 h.

**Data collection and structure determination.** The soaked crystals were cryoprotected by snap-freezing in liquid nitrogen with a well solution containing ethylene glycol (20%) and the tested compound (1.0 mM). Diffraction data were collected on beamlines BL17U1 and BL19U1 at Shanghai Synchrotron Radiation Facility (SSRF). The data were processed by using HKL3000.^7^ The structure was determined by molecular replacement with the T2R−TTL complex structure (PDB ID: 4O2B) as a searching model in Phenix.^8^ The resulting model was built in COOT^9^ and refined in Phenix.^8^ Coordinates have been deposited in the Protein Data Bank with identification codes 7CPD [T2R−TTL−(+)-**8**] and 7CPQ [T2R−TTL−(+)-**9**]. For data collection and refinement statistics, see Table S1.

**RNA extraction, reverse transcription, and quantitative real-time PCR.** HeLa cells were seeded in a 6-well culture plate at a density of 4.0 × 10^5^ per well and treated with (+)-**8** (0.50 *μ*M), (+)-**9** (0.50 *μ*M), nocodazole (0.10 *μ*M), and vinblastine (0.050 *μ*M), respectively, for 6 h. DMSO was used as the vehicle for the compound. Total RNA was extracted by using TRIzol Reagent (Invitrogen). Reverse transcription was performed by using PrimeScript RT Reagent Kit with gDNA Eraser (Takara). Quantitative real-time PCR (qRT-PCR) was performed with QuantStudio Real-Time PCR System (Thermo Fisher Scientific). The mRNA levels of β-actin were used as the reference for normalization. The following sense (S) and antisense (AS) primers were used for qRT-PCR.

CENP-A (S): 5’-CTTCCTCCCATCAACACAGTCG-3’.

CENP-A (AS): 5’-TGCTTCTGCTGCCTCTTGTAGG-3’.

β-actin (S): 5’-GAGCGGGAAATCGTGCGTGACATT-3’.

β-actin (AS): 5’-GATGGAGTTGAAGGTAGTTTCGTG-3’.

**Statistical analysis.** Statistical analyses were performed with GraphPad Prism 5.0. The two-tailed unpaired Student's *t*-test was used to calculate differences between the two groups. Differences were considered statistically significant at **P* < 0.05, ***P* < 0.01, ****P* < 0.001, and *****P* < 0.0001. All data shown were presented as mean ± s.e.m. and represented at least three independent experiments.

**II	Figures and Tables**

**Figure S1. Cell cycle-arresting effect of JP18 (1) and its analogues.** HeLa cells were treated with the indicated compounds at the indicated concentrations for 8 h and then stained with propidium iodide (PI). DMSO was used as a control for the compounds. Cell cycle distribution was analyzed by flow cytometry.

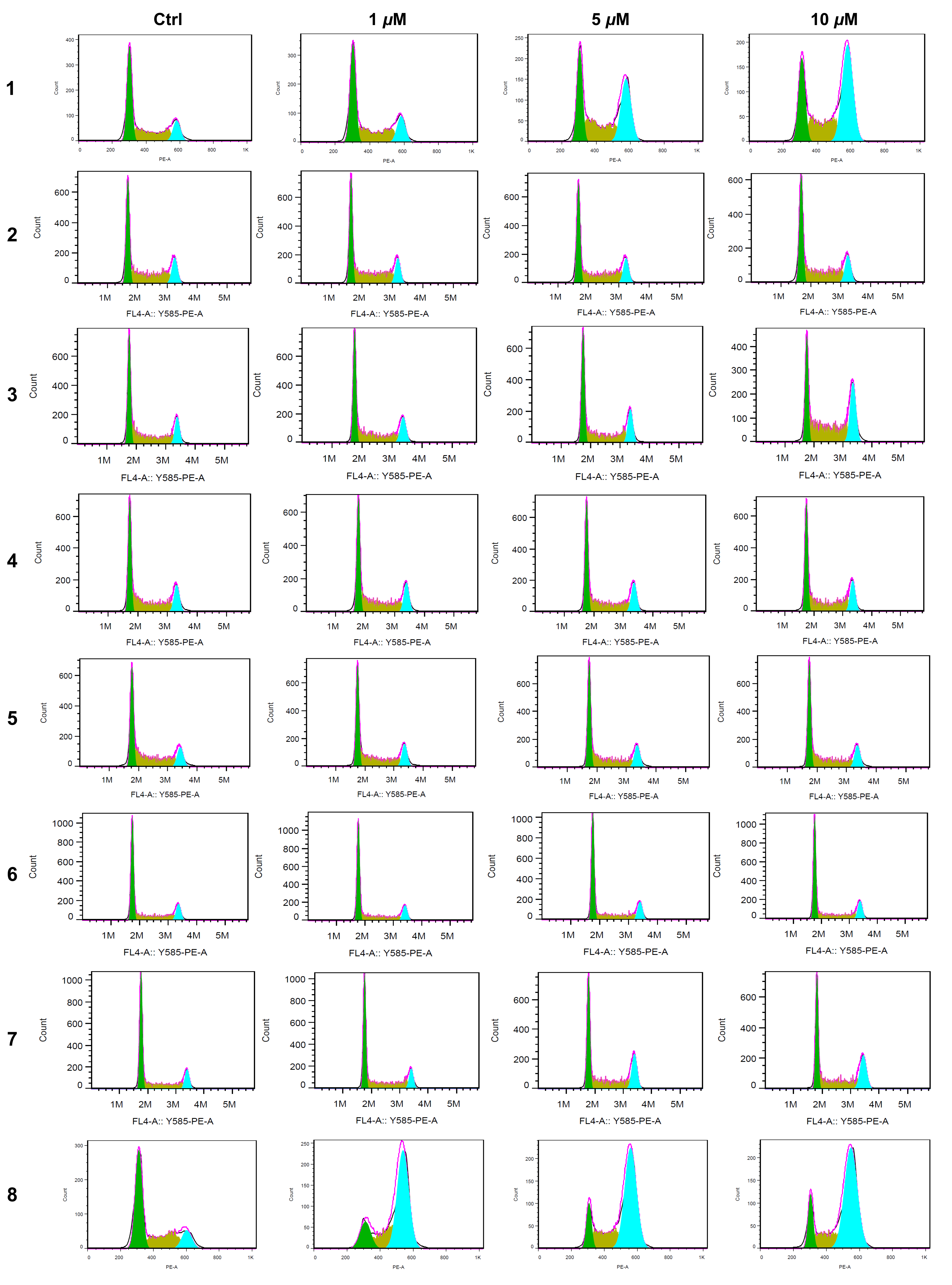

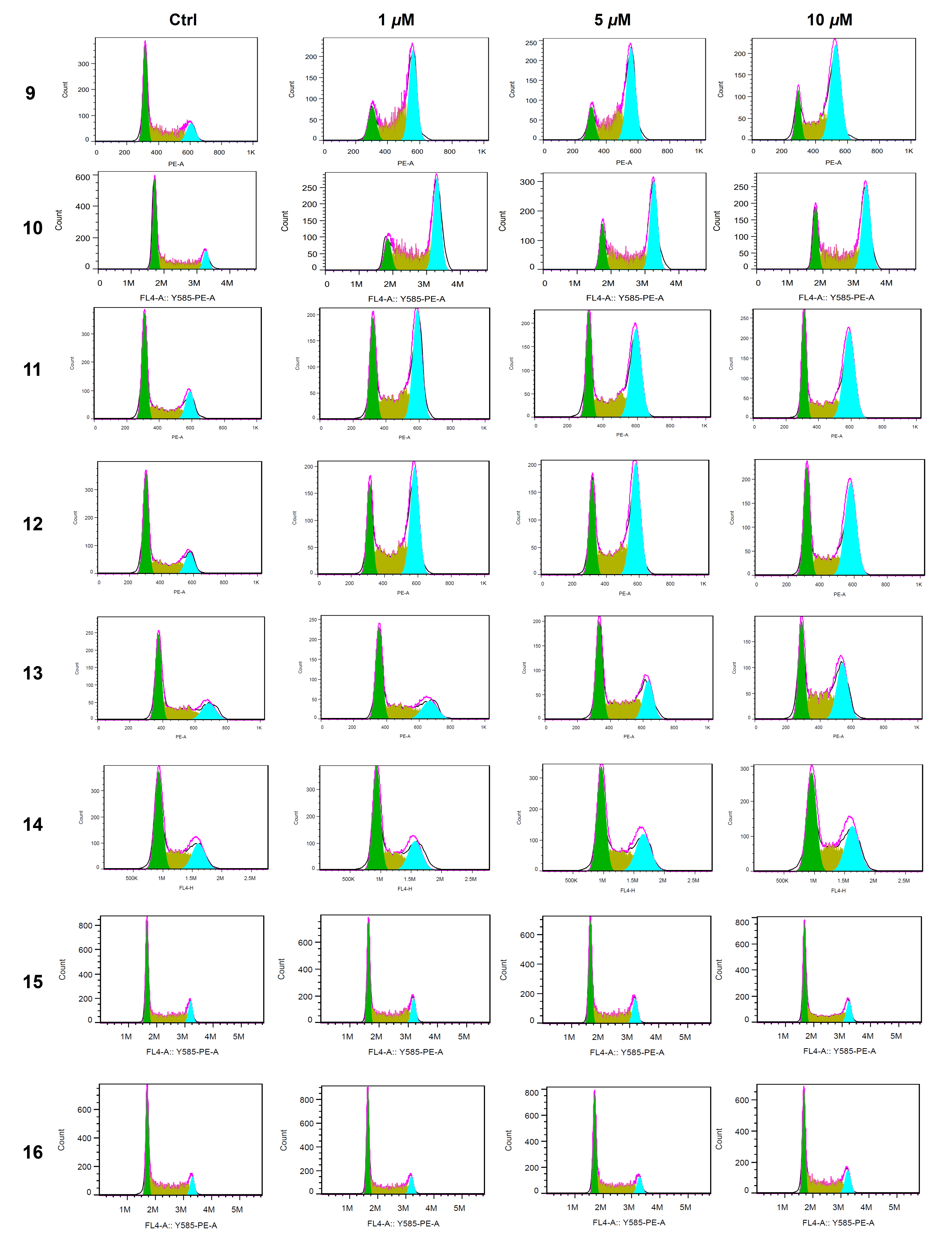

**Figure S2. Compounds 8 and 9 exhibit stronger G2/M phase-arresting activity than compound 1.** (A) Compounds **8** and **9** induced cell cycle arrest in the G2/M phase in a dose-dependent manner at low concentrations. HeLa cells were treated with compounds **1**, **8**, and **9** at the indicated concentrations for 8 h. (B) Compounds **8** and **9** induced cell cycle arrest in the G2/M phase at 0.50 *μ*M in a time-dependent manner. HeLa cells were treated with compounds **1**, **8**, and **9** at 0.50 *μ*M for the indicated durations. DMSO was used as a control for the compounds. Bar graphs show the percentages of cells in the G2/M (green), S (yellow), and G1 (orange) phases.

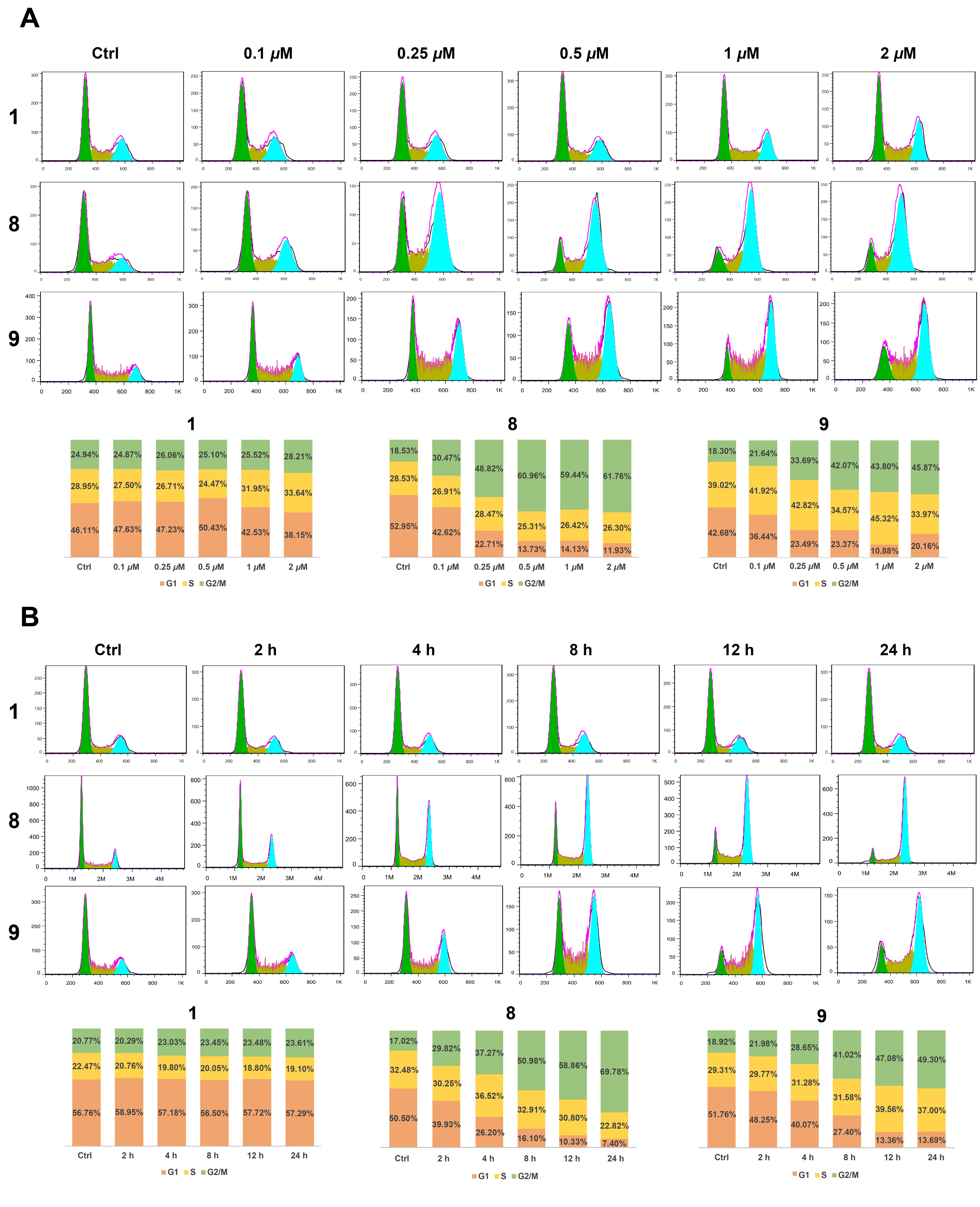

**Figure S3. Curves for determining the IC_50_ of compounds 1, 8 and 9 in HeLa cells.** HeLa cells were treated with the indicated compounds at concentrations ranging from 0 to 10 *μ*M for 24 h. Cell viability was measured by using CCK-8. Data were analyzed with GraphPad Prism 5.0.

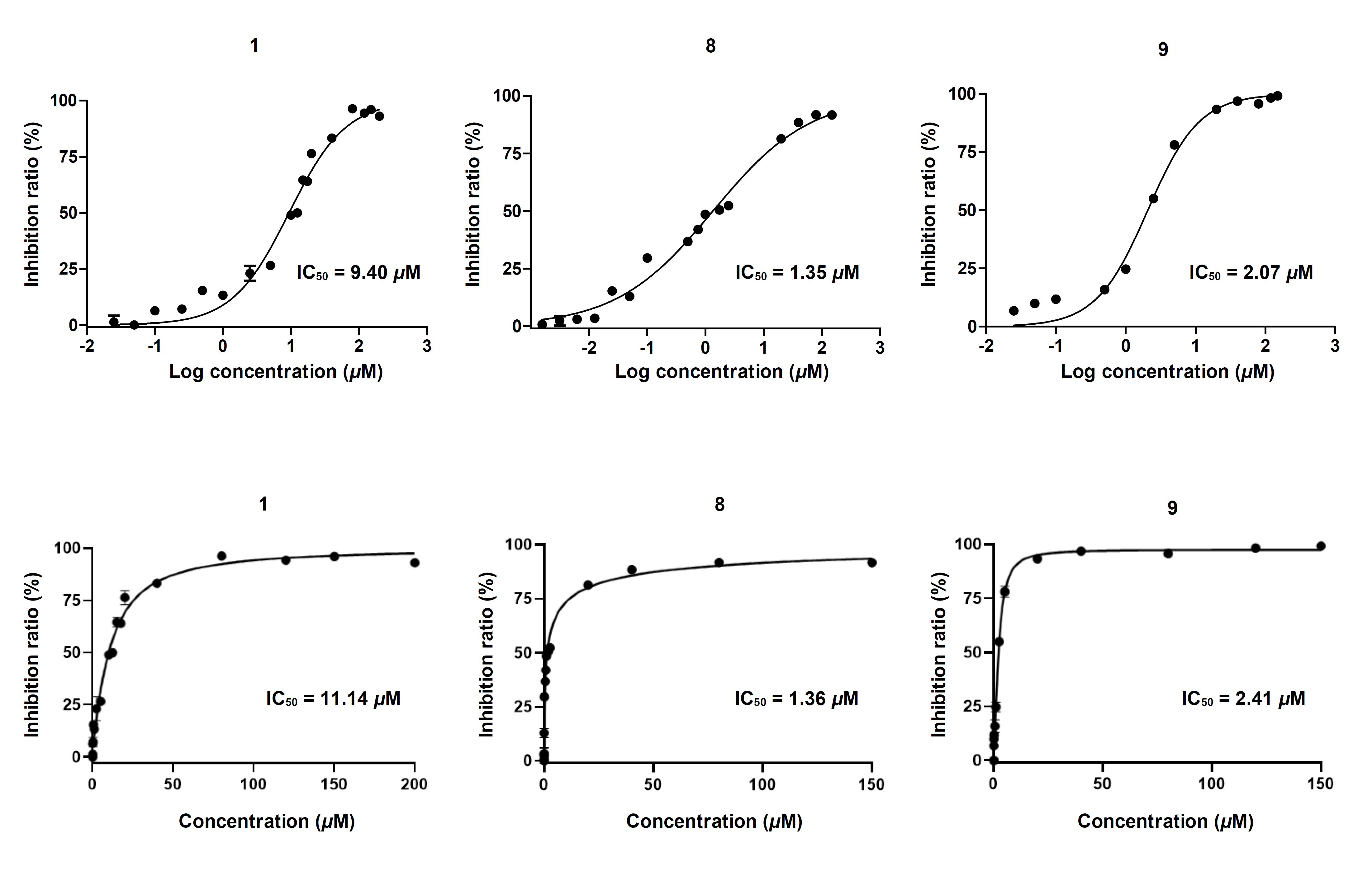

**Figure S4. Determination of the enantiomeric forms of compounds 8 and 9 responsible for the cell cycle-arresting activity.** (A) Comparison of the cell cycle-arresting effect of (+)-**8** and (−)-**8**. HeLa cells were treated with the indicated compound at the indicated concentrations for 8 h. (B) Comparison of the cell cycle arresting effect of (+)-**9** and (−)-**9**. HeLa cells were treated with the indicated compound at the indicated concentrations for 8 h. DMSO was used as a control for the compounds.

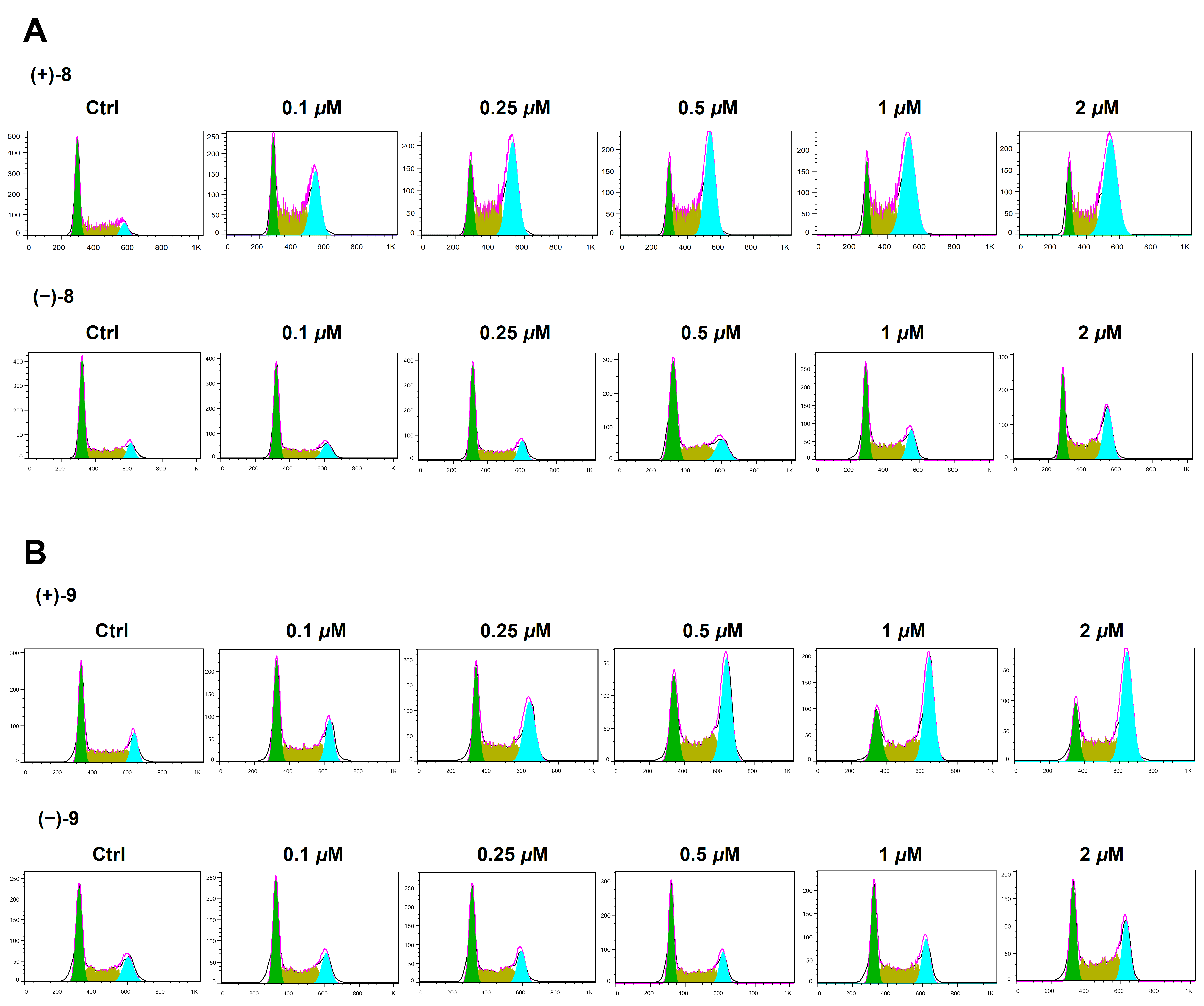

**Figure S5. Effect of compounds (+)-8 and (+)-9 on markers of apoptosis and DNA damage.** (A, B) Immunoblot analysis of cleaved PARP1, Bcl-2, and γ-H2AX in HeLa cells treated with (+)-**8** and (+)-**9**, respectively, at the indicated concentrations for 24 h. (C, D) Immunoblot analysis of cleaved PARP1, Bcl-2, and γ-H2AX in HeLa cells treated with (+)-**8** and (+)-**9**, respectively, at 0.50 *μ*M for the indicated durations. DMSO was used as a control for the compounds. GAPDH was used as a loading control in immunoblotting.

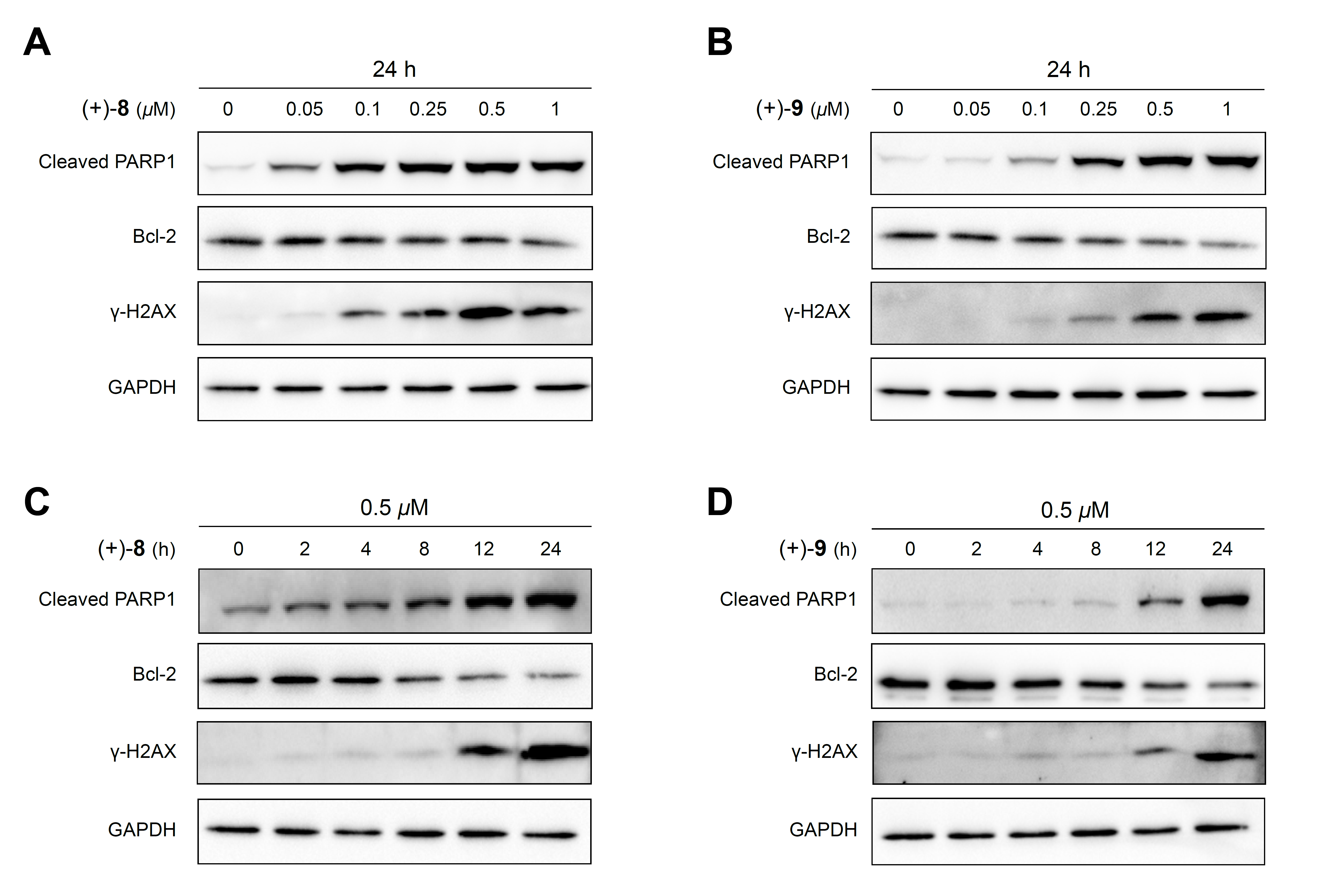

**Figure S6.** Overall structure of the T2R−TTL−(+)-**9** complex. α-Tubulin (light slate blue), β-tubulin (green), RB3-SLD (plum), and TTL (gold) are shown in cartoon representation. Compound (+)-**9**, GDP, and GTP are shown in sphere representation.

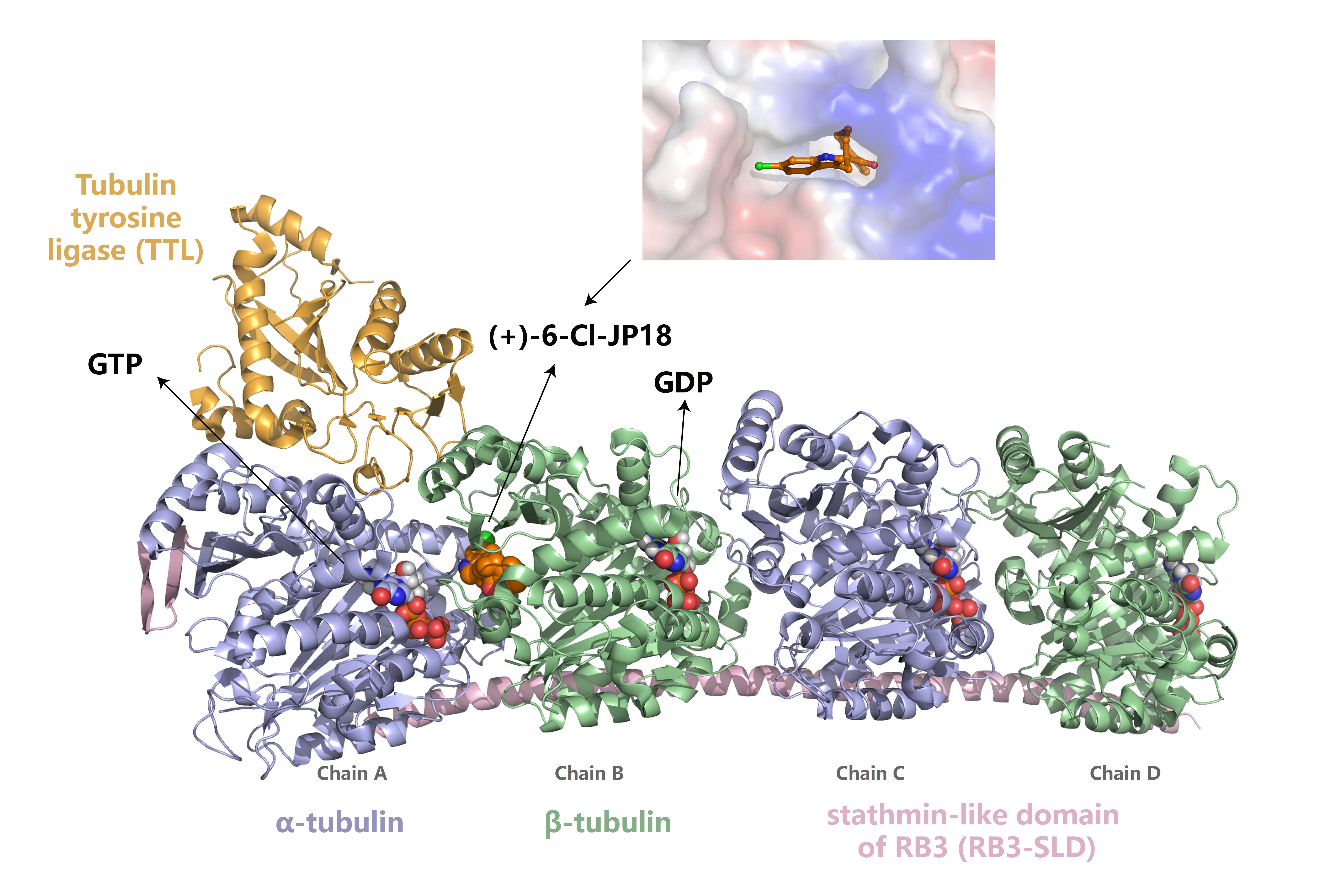

**Figure S7. Analysis of the post-translational regulation of CENP-A.** (A) Immunoblot analysis of CENP-A in HeLa cells treated with NH_4_Cl (40 mM) for 3 h. (B) Immunoblot analysis of CENP-A in HeLa cells treated with CQ (30 *μ*M) for 3 h. LC3B was used as a positive control. (C) Immunoblot analysis of CENP-A in HeLa cells transfected with Myc-NEDD8. (D) Immunoblot analysis of CENP-A in HeLa cells transfected with Myc-SUMO1. (E) Comparison of the CENP-A levels in HeLa cells treated with nocodazole (0.10 *μ*M) and co-treated with nocodazole (0.10 *μ*M) and MG-132 (50 *μ*M) for the indicated durations by immunoblotting. *4 h. **6 h.

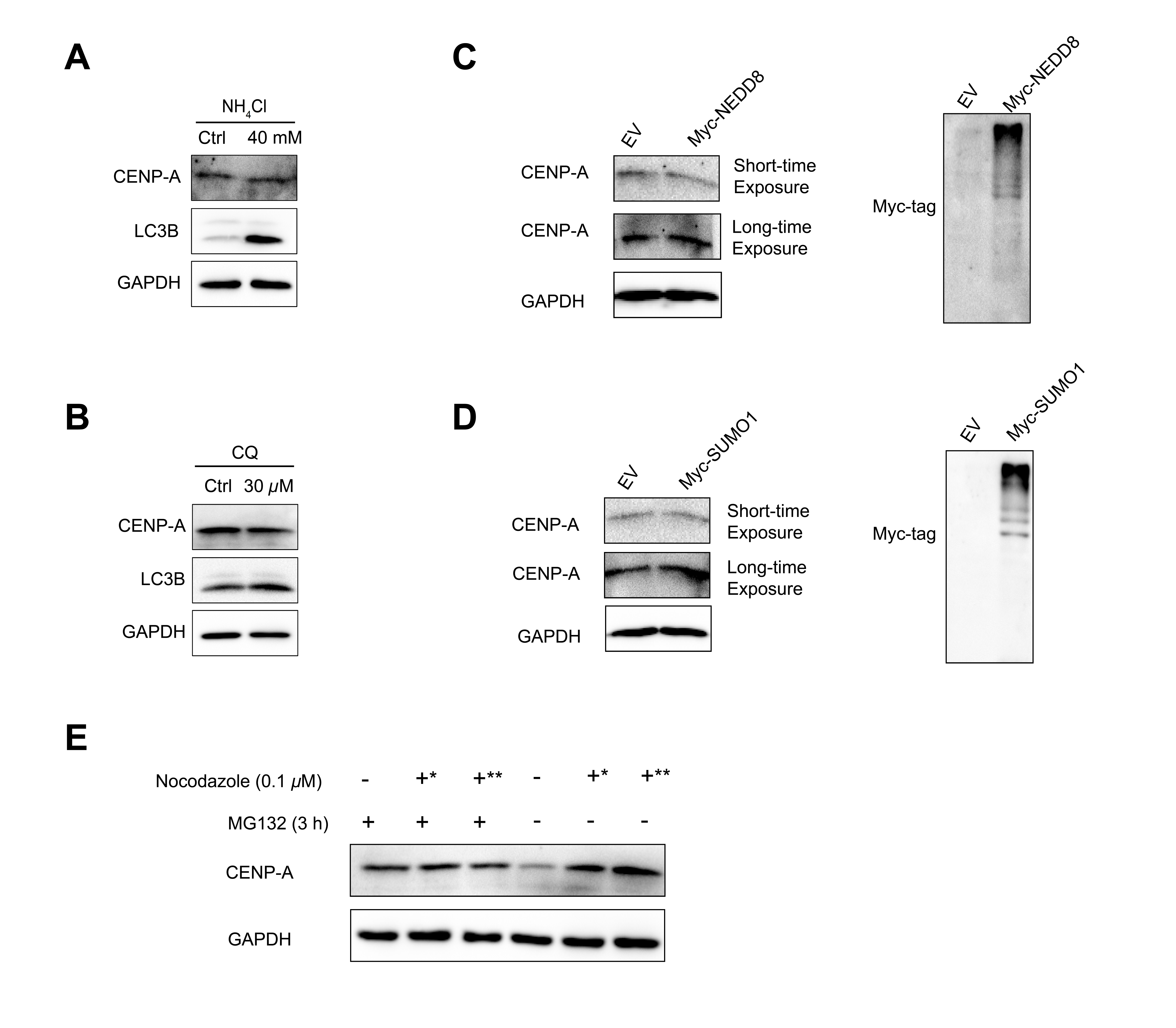

**Figure S8. Compound (+)-8 and paclitaxel downregulate Cdh1 and upregulate CENP-A in various human cells.** (A) Immunoblot analysis of CENP-A, Cdh1, and Cdc20 in Hep G2 cells treated with (+)-**8** (0.50 *μ*M) for the indicated durations. (B) Immunoblot analysis of CENP-A, Cdh1, and Cdc20 in A549 cells treated with (+)-**8** (0.50 *μ*M) for the indicated durations. (C) Immunoblot analysis of CENP-A, Cdh1, and Cdc20 in L-02 cells treated with (+)-**8** (0.50 *μ*M) for the indicated durations. (D) Immunoblot analysis of CENP-A, Cdh1, and Cdc20 in Hep G2 cells treated with paclitaxel (0.20 *μ*M) for the indicated durations. (E) Immunoblot analysis of CENP-A, Cdh1, and Cdc20 in A549 cells treated with paclitaxel (0.20 *μ*M) for the indicated durations. (F) Immunoblot analysis of CENP-A, Cdh1, and Cdc20 in L-02 cells treated with paclitaxel (0.20 *μ*M) for the indicated durations. DMSO was used as a control for the compounds. GAPDH was used as a loading control in immunoblotting.

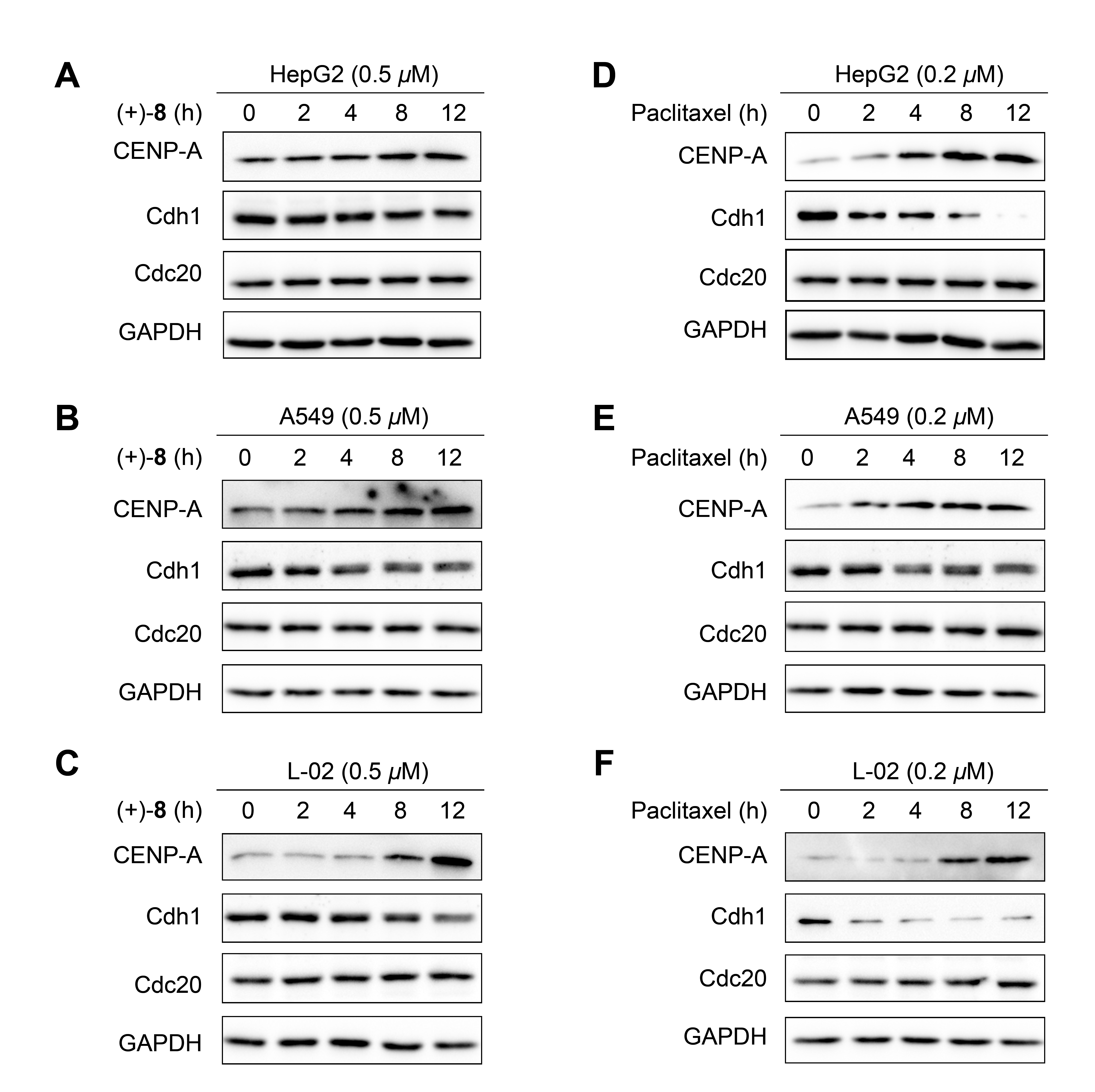

**Figure S9. Compound (+)-8 downregulates Cdh1 at the post-translational level.** The Cdh1 levels in HeLa cells treated with CHX (100 *μ*g/mL) alone and co-treated with CHX (100 *μ*g/mL) and (+)-**8** (0.50 *μ*M) for the indicated durations were analyzed by immunoblotting. DMSO was used as a control for the compound. GAPDH was used as a loading control in immunoblotting.

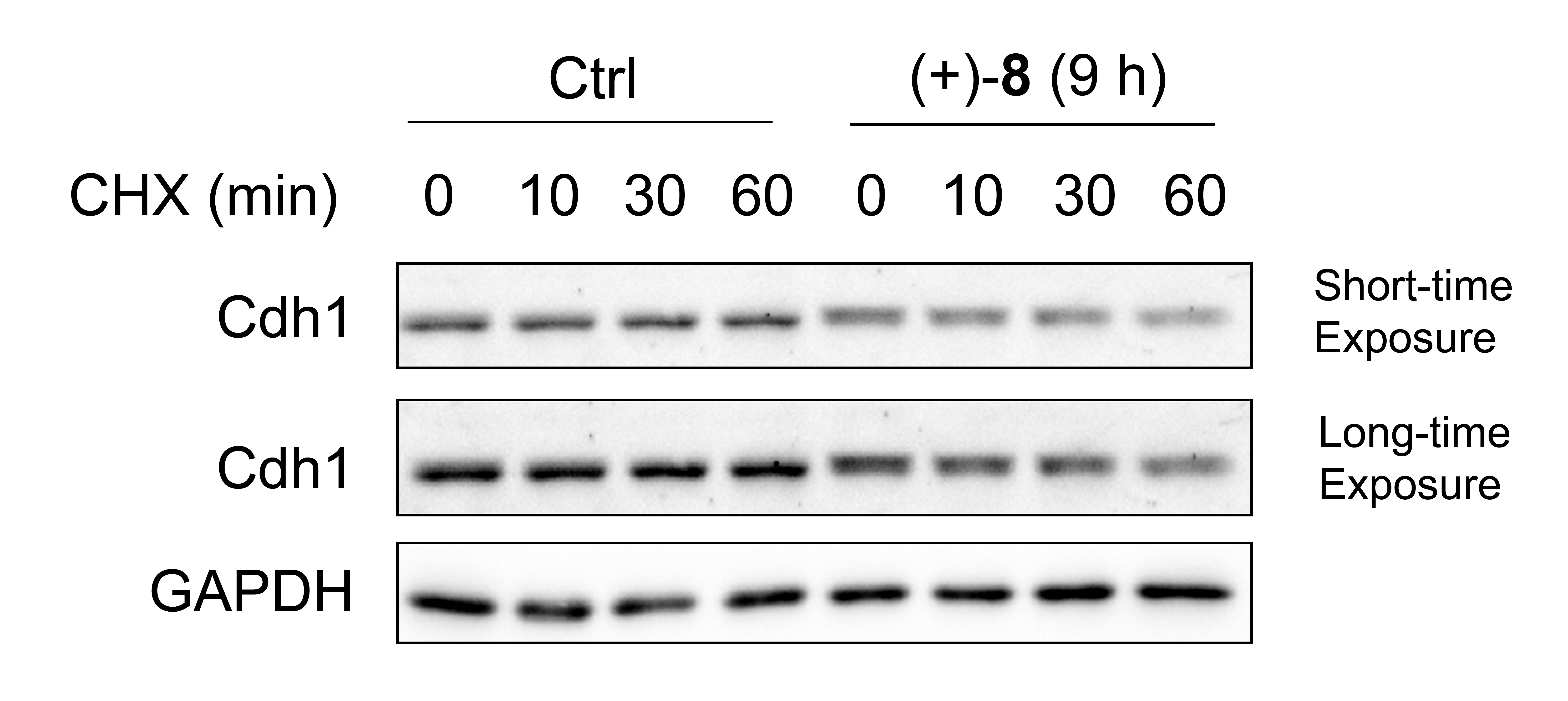

**Figure S10. Cdc20 knockdown does not affect CENP-A levels.** (A) Immunoblot analysis of CENP-A in HeLa cells transfected with siRNA targeting Cdc20 mRNA. (B) Quantitative analysis of the immunoblotting data obtained from the above experiment. Data are representative of three independent biological replicates and are presented as mean ± s.e.m. NS = not significant (significance level: α = 0.05; *n* = 6, two-tailed Student’s *t*-test).

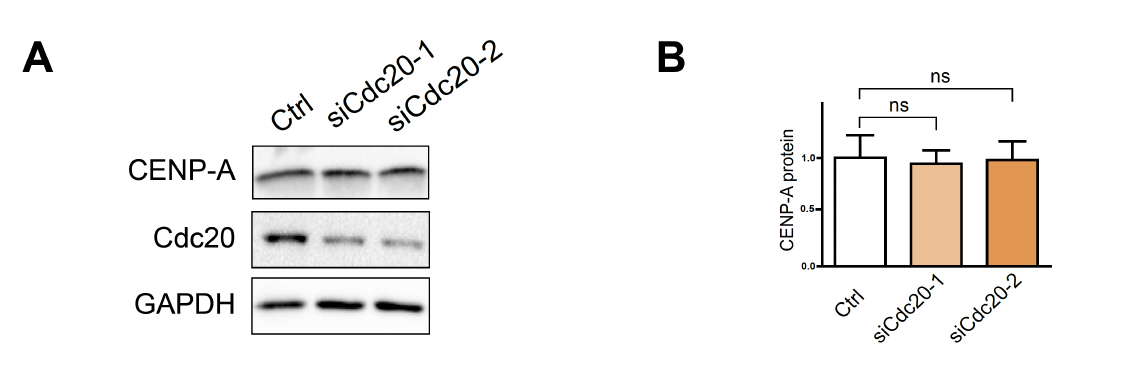

**Figure S11. Cdh1 knockdown increases CENP-A in various human cells.** (A–E) Immunoblot analysis of CENP-A in Hep G2, A549, MDA-MB-231, U251, and L-02 cells transfected with siRNA targeting Cdh1 mRNA, respectively. Two different siRNA oligos were employed independently. siRNA targeting a non-relevant mRNA was used as a negative control. GAPDH was used as a loading control in immunoblotting.

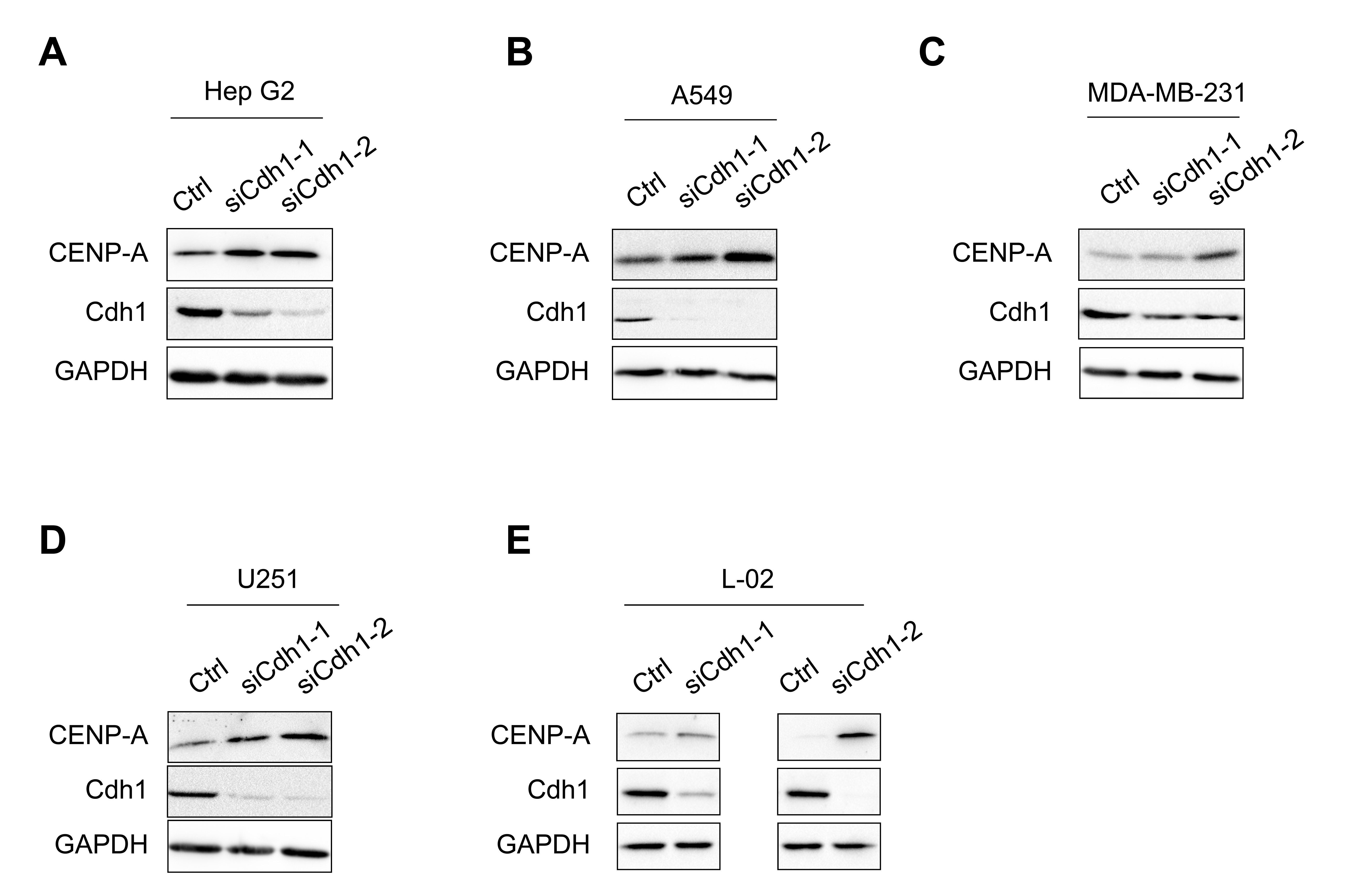

**Figure S12. Combination of Cdh1 knockdown and MTA treatment enhances CENP-A accumulation compared to MTA treatment alone.** (A) Immunoblot analysis of CENP-A in HeLa cells transfected with siRNA targeting Cdh1 mRNA and then treated with (+)-**8** (0.50 *μ*M) for 12 h. (B) Immunoblot analysis of CENP-A in HeLa cells transfected with siRNA targeting Cdh1 mRNA and then treated with paclitaxel (0.20 *μ*M) for 12 h. (C) Immunoblot analysis of CENP-A in HeLa cells transfected with siRNA targeting DCAF11 mRNA and then treated with (+)-**8** (0.50 *μ*M) for 12 h. (D) Immunoblot analysis of CENP-A in HeLa cells transfected with siRNA targeting DCAF11 mRNA and then treated with paclitaxel (0.20 *μ*M) for 12 h. Two different siRNA oligos were employed independently. siRNA targeting a non-relevant mRNA was used as a negative control. GAPDH was used as a loading control in immunoblotting.

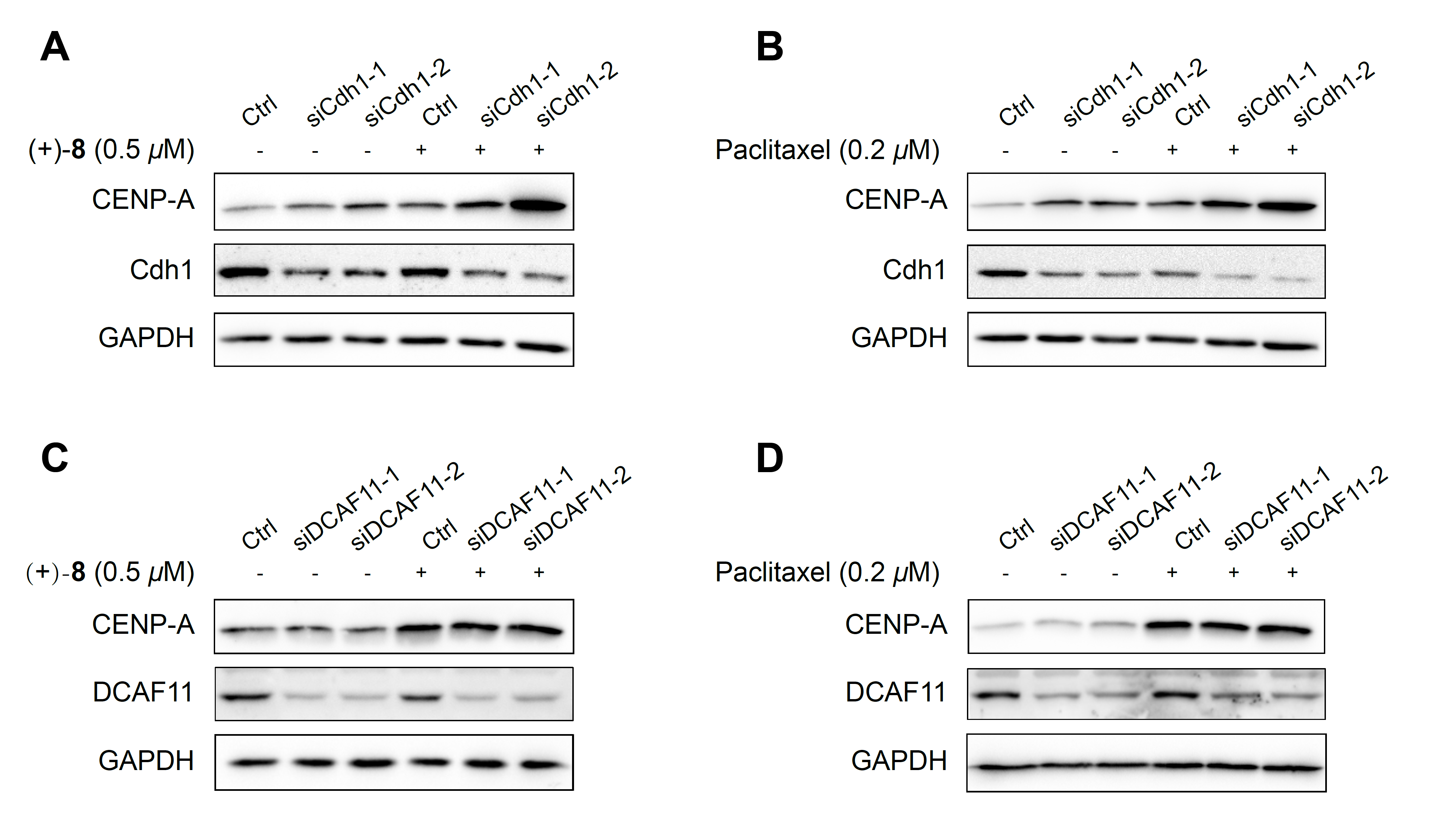

**Figure S13. Combination of APC2 knockdown and MTA treatment enhances CENP-A accumulation compared to MTA treatment alone.** (A) Immunoblot analysis of CENP-A in HeLa cells transfected with siRNA targeting APC2 mRNA and then treated with (+)-**8** (0.50 *μ*M) for 12 h. (B) Immunoblot analysis of CENP-A in HeLa cells transfected with siRNA targeting APC2 mRNA and then treated with paclitaxel (0.20 *μ*M) for 12 h. Two different siRNA oligos were employed independently. siRNA targeting a non-relevant mRNA was used as a negative control. GAPDH was used as a loading control in immunoblotting.

**Figure S14. Compound (+)-8 and paclitaxel do not affect Cullin 4A or DCAF11 levels.** (A) Immunoblot analysis of Cullin 4A and DCAF11 in HeLa cells treated with (+)-**8** (0.50 *μ*M) for the indicated durations. (B) Immunoblot analysis of Cullin 4A and DCAF11 in HeLa cells treated with paclitaxel (0.20 *μ*M) for the indicated durations.

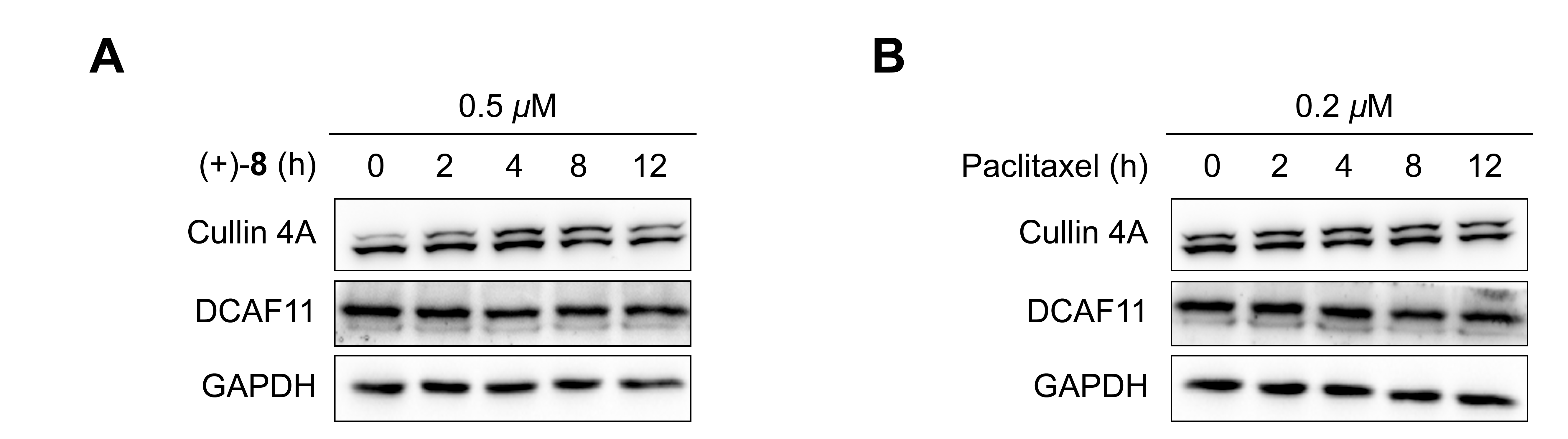

**Table S1. Data collection and refinement statistics.**

|  | T2R−TTL−(+)-**8** complex | T2R−TTL−(+)-**9** complex |
| --- | --- | --- |
| Wavelength | 0.9785 | 0.9875 |
| Resolution range | 49.89–2.506 (2.596–2.506) | 48.06–2.595 (2.688–2.595) |
| Space group | P 21 21 21 | P 21 21 21 |
| Unit cell | 104.782 156.552 182.591  90 90 90 | 104.877 156.616 182.624  90 90 90 |
| Total reflections | 1335935 (107124) | 1201389 (76058) |
| Unique reflections | 102800 (8435) | 92838 (7549) |
| Redundancy | 13.0 (12.7) | 12.9 (11.4) |
| Completeness (%) | 99.4 (99.4) | 99.4 (99.7) |
| Mean I/sigma (I) | 26.40 (2.67) | 23.00 (2.00) |
| Wilson B-factor | 35.09 | 40.20 |
| R-merge | 0.089 (0.987) | 0.206 (2.023) |
| R-meas | 0.093 (1.028) | 0.215 (2.116) |
| CC1/2 | 0.995 (0.813) | 0.987 (0.708) |
| Reflections used in refinement | 100497 (8434) | 90558 (7550) |
| Reflections used for R-free | 5063 (434) | 4654 (380) |
| R-work | 0.1829 (0.2288) | 0.2193 (0.2954) |
| R-free | 0.2401 (0.3027) | 0.2664 (0.3548) |
| Number of non-hydrogen atoms | 17898 | 17602 |
| Macromolecules | 17326 | 17326 |
| Ligands | 191 | 190 |
| Protein residues | 2162 | 2162 |
| RMS (bonds) | 0.011 | 0.012 |
| RMS (angles) | 0.98 | 0.92 |
| Ramachandran favored (%) | 95 | 95 |
| Ramachandran allowed (%) | 4.3 | 4.6 |
| Ramachandran outliers (%) | 0.6 | 0.78 |
| Rotamer outliers (%) | 3.7 | 4.2 |
| Clashscore | 17.46 | 14.44 |
| Average B-factor | 51.47 | 46.77 |
| Macromolecules | 51.65 | 46.87 |
| Ligands | 45.20 | 40.76 |
| Solvent | 46.35 | 39.53 |
| Number of TLS groups | 1 | 1 |

Statistics for the highest-resolution shell are shown in parentheses.

**III	References**

1. X. Xiong, D. Zhang, J. Li, Y. Sun, S. Zhou, M. Yang, H. Shao, A. Li, Synthesis of indole terpenoid mimics through a functionality-tolerated Eu(fod)_3_-catalyzed conjugate addition. *Chem. Asian J.* 10, 869–872 (2015).

2. Y. Sun, P. Chen, D. Zhang, M. Baunach, C. Hertweck, A. Li, Bioinspired total synthesis of sespenine. *Angew. Chem. Int. Ed.* 53, 9012–9016 (2014).

3. J. Pei, S. Zhou, F. Yang, Y. Sun, A. Li, W. D. Zhang, W. He, Identification and mechanistic studies of a cell cycle regulator JP18 from a library of synthetic indole terpenoid mimics. *Chem. Asian J.* 11, 2715–2718 (2016).

4. B. Lomenick, R. Hao, N. Jonai, R. M. Chin, M. Aghajan, S. Warburton, J. Wang, R. P. Wu, F. Gomez, J. A. Loo, J. A. Wohlschlegel, T. M. Vondriska, J. Pelletier, H. R. Herschman, J. Clardy, C. F. Clarke, J. Huang, Target identification using drug affinity responsive target stability (DARTS). *Proc. Natl. Acad. Sci. USA* 106, 21984–21989 (2009).

5. M. Y. Pai, B. Lomenick, H. Hwang, R. Schiestl, W. McBride, J. A. Loo, J. Huang, Drug affinity responsive target stability (DARTS) for small-molecule target identification. *Methods Mol. Biol*. 1263, 287–298 (2015).

6. A. E. Prota, M. M. Magiera, M. Kuijpers, K. Bargsten, D. Frey, M. Wieser, R. Jaussi, C. C. Hoogenraad, R. A. Kammerer, C. Janke, M. O. Steinmetz, Structural basis of tubulin tyrosination by tubulin tyrosine ligase. *J. Cell Biol*. 200, 259–270 (2013).

7. W. Minor, M. Cymborowski, Z. Otwinowski, M. Chruszcz, HKL-3000: the integration of data reduction and structure solution — from diffraction images to an initial model in minutes. *Acta Crystallogr. D* 62, 859–866 (2006).

8. P. D. Adams, P. V. Afonine, G. Bunkoczi, V. B. Chen, I. W. Davis, N. Echols, J. J. Headd, L. W. Hung, G. J. Kapral, R. W. Grosse-Kunstleve, A. J. McCoy, N. W. Moriarty, R. Oeffner, R. J. Read, D. C. Richardson, J. S. Richardson, T. C. Terwilliger, P. H. Zwart, PHENIX: a comprehensive Python-based system for macromolecular structure solution. *Acta Crystallogr. D* 66, 213–221 (2010).

9. P. Emsley, B. Lohkamp, W. G. Scott, K. Cowtan, Features and development of Coot. *Acta Crystallogr. D* 66, 486–501 (2010).
